# Minimum complexity drives regulatory logic in Boolean models of living systems

**DOI:** 10.1101/2021.09.20.461164

**Authors:** Ajay Subbaroyan, Olivier C. Martin, Areejit Samal

## Abstract

The properties of random Boolean networks as models of gene regulation have been investigated extensively by the statistical physics community. In the past two decades, there has been a dramatic increase in the reconstruction and analysis of Boolean models of biological networks. In such models, neither network topology nor Boolean functions (or logical update rules) should be expected to be random. In this contribution, we focus on *biologically meaningful* types of Boolean functions, and perform a systematic study of their preponderance in gene regulatory networks. By applying the *k*[*P*] classification based on number of inputs *k* and bias *P* of functions, we find that most Boolean functions astonishingly have odd bias in a reference biological dataset of 2687 functions compiled from published models. Subsequently, we are able to explain this observation along with the enrichment of read-once functions (RoFs) and its subset, nested canalyzing functions (NCFs), in the reference dataset in terms of two complexity measures: *Boolean complexity* based on string lengths in formal logic which is yet unexplored in the biological context, and the *average sensitivity*. Minimizing the Boolean complexity naturally sifts out a subset of odd-biased Boolean functions which happen to be the RoFs. Finally, we provide an *analytical* proof that NCFs minimize not only the Boolean complexity, but also the average sensitivity in their *k*[*P*] set.

## I. INTRODUCTION

Cells are the building blocks of all living organisms and their decision-making is tightly controlled by complex and intricate gene regulatory networks [1]. The statistical physics community in particular have contributed immensely to the deeper understanding of the structure and dynamics of these complex biological networks [2–9]. Discrete logic modeling was introduced by Stuart Kauffman [10, 11] and René Thomas [12, 13] as a framework to study the dynamics of gene regulatory networks. The Boolean framework though qualitative in approach (in that all biological entities are approximated to switches being “on” or “off”), can remarkably reproduce the expected steady states of biological systems [14, 15]. Kauffman studied ensembles of Boolean networks drawn from prior distributions, specifically, NK-Boolean networks [10]. Since then, the NK-Boolean networks and the nature of their attractors under various updating schemes have been studied quite extensively [2, 16, 17]. Notably, the Boolean functions (BFs) or logic rules are chosen at random while studying the dynamics of these NK-Boolean networks.

Advances in large-scale data generation has led to the construction of biological networks in many organisms [5, 18]. Extensive studies of these real networks has revealed that their structure is very different from being random [5, 6, 9]. Moreover, Boolean dynamical models for several biological systems [19–25] have been constructed in the last two decades. It will be important to characterize the properties associated with the logical update rules or BFs in these real models that distinguish them from randomly chosen functions. In previous studies [2, 26, 27], one property that has been used to characterize the BFs is their bias, which is the probability of a BF to have an output value 1. Though the bias is very useful in characterizing ensembles of Boolean models, the normalized quantity may camouflage key information associated with certain types of BFs. By using instead the unnormalized bias *P*, also known as activity [26] or Hamming weight, we elucidate important properties of certain types of BFs. Specifically, we employ the *k*[*P*] classification scheme by Feldman [28] wherein *k* is the number of inputs to characterize BFs.

Multiple biologically meaningful types of BFs have been introduced previously in the literature such as the effective functions (EFs) [29], unate functions (UFs) [30], canalyzing functions (CFs) [2, 31] and nested canalyzing functions (NCFs) [21, 32, 33]. Due to possible overlap among these types of BFs, a solitary exploration of any type is insufficient. Hence, we systematically study the different types of biologically meaningful functions and their relationships in both the complete space of 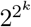 BFs and a reference biological dataset of 2687 BFs compiled from 88 published models. Furthermore, we look at these different types of biologically meaningful functions through the lens of *k*[*P*] classification [28]. This led to multiple theoretical conjectures on relevant properties of these BF types for which we are able to provide analytical proofs.

Kauffman [2] had proposed that “chemical simplicity” could be a reason for the widespread use of CFs in nature. We borrow concepts from the computer science literature to quantify the notion of simplicity (or complexity) of a BF, and thereafter, perform a thorough evaluation of the biologically meaningful types of functions from the perspective of complexity. The two measures of complexity which we explore are the Boolean complexity [28] and the average sensitivity [26, 34]. We show that RoFs which constitute all logic rules with minimal Boolean complexity are highly over-represented in the biological data. Further, we provide an analytical proof that NCFs, which are a subset of RoFs, minimize not only the Boolean complexity but also the average sensitivity across all BFs in their *k*[*P*] set. Our result that NCFs are minimally complex in terms of two complexity measures is a likely explanation for their prevalence in the biological data. In a nutshell, our exploration of the two complexity measures in the biological data of 2687 BFs compiled from published models confirms the proposition made by Kauffman on preference for simplicity among the choices of logic rules in gene regulatory networks.

## II. BACKGROUND

### A. Boolean models of biological networks

A Boolean model of a biological system consists of a network of *N* nodes and *L* edges wherein the nodes correspond to components such as genes or proteins and (directed) edges capture the regulation of one node by another node [2, 10–12]. Let us label each node of the network by an integer *i* (*i* = 1,…, *N*) and denote the state of each node *i* by a Boolean variable *x_i_* ∈ {0, 1}. The state *x_i_* of a node *i* in the Boolean model is determined by: (a) the values of its *k_i_* inputs, coming from the *k_i_* nodes from which it has incoming links, and (b) a logical update rule or *Boolean function f_i_* that specifies how *x_i_* changes in time or *is updated* given those *k_i_* inputs. Thus, every node *i* in the network with at least one input is associated with a Boolean function (BF) which governs its dynamics. There are many ways to update the list of *x_i_* values across the network. One is the *synchronous* update procedure [2] whereby all nodes of the network are simultaneously updated. When the nodes are not all updated in that way, the dynamics is said to be *asynchronous* [12, 13]. In this contribution, such different update procedures can be ignored because we are primarily concerned with characterizing the types of BFs that are abundant in Boolean models of biological networks.

### B. Representations, Bias and Parity of Boolean functions

Let *f* = *f*(*x*_1_, *x*_2_,…, *x_k_*) be a BF of *k* input variables where each variable *x_i_* ∈ {0, 1}. A BF maps 2^*k*^ different possibilities for the *k* input variables to output values 0 or 1, i.e., *f*: {0, 1}^*k*^ → {0, 1}.

#### Truth table and associated ordered binary vector

A BF *f* with *k* inputs can be captured in the form of a truth table with 2^*k*^ rows, each corresponding to a possible state of the set of *k* input variables. In our convention the last entry of each row gives the output value for the corresponding state of the input variables (Fig. 1(a)). Thus, a BF can be stored as a binary vector of size 2^*k*^, where each element of the vector corresponds to the output value of the corresponding row of the truth table (Fig. 1(b)). A BF can also be encoded as the integer which is the decimal equivalent of the binary vector of size 2^*k*^. Since the output value for each of the 2^*k*^ different states of the *k* input variables can take either of two values, 0 or 1, the number of possible BFs *f* with *k* inputs is 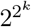. Thus, the number of possible BFs *f* blows up quickly with increasing *k* [2], e.g., there are over 10^9^ BFs with *k* = 5 inputs (Table S1, Figure S1).

**FIG. 1.**
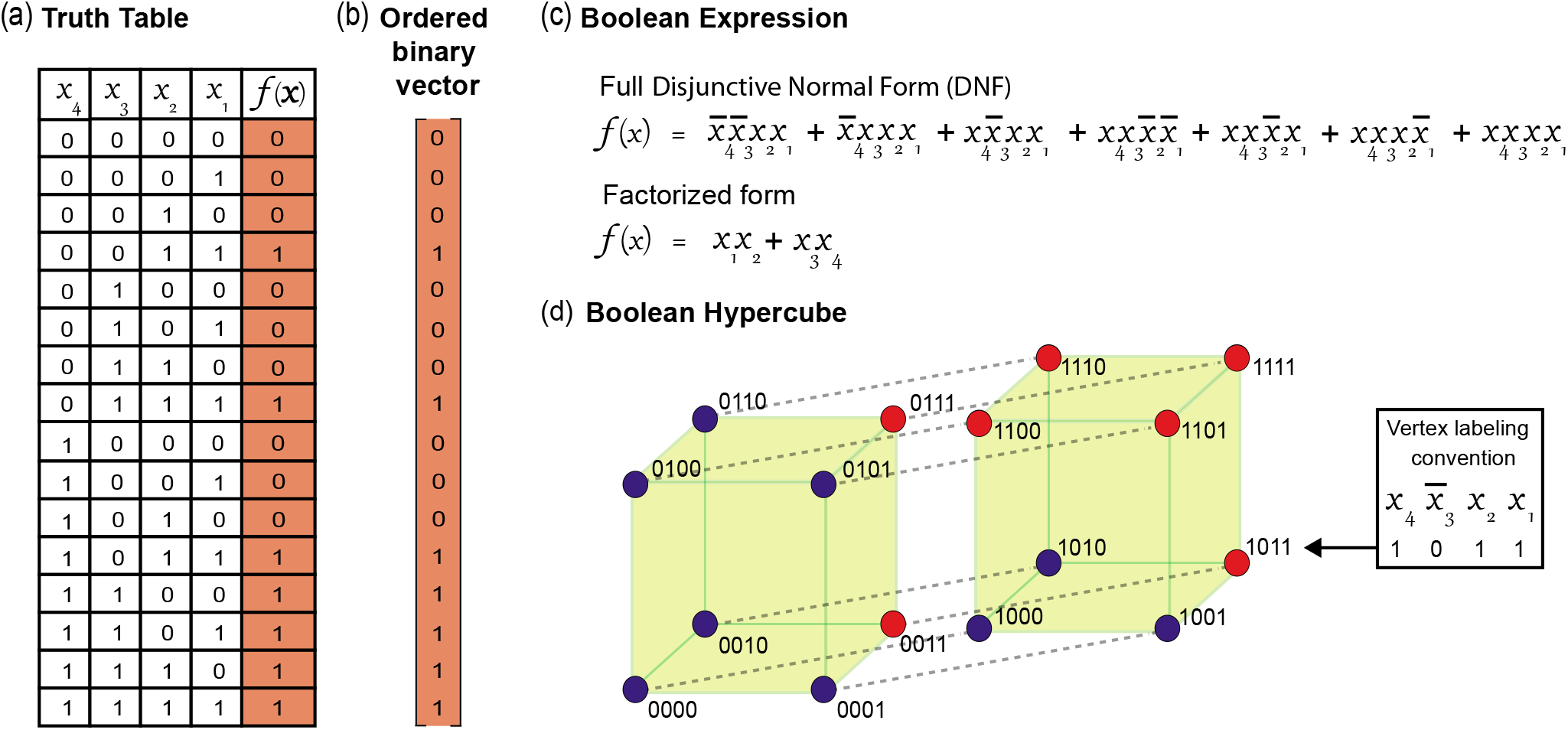
Illustration of four different representations of Boolean functions (BFs) using an example. **(a)** Truth table. **(b)** Ordered binary vector, which is just the output column of the truth table, but now taken as a *k*-dimensional vector. **(c)** Boolean expression. Multiple types of Boolean expressions are possible. We have shown for illustration the BF in the truth table in its Full Disjunctive Normal Form (DNF) and it’s minimum equivalent expression. In the Full DNF, each row of the truth table whose output value is 1 is represented as a product of *k* literals, each one corresponding to one input bit. If an input bit is 0, it is represented by a negated literal, otherwise, by a positive one. **(d)** Colored Boolean Hypercube. Each vertex corresponds to one row of the truth table. The position of the vertex is defined by the inputs and the color of the vertex by the output corresponding to that input. The color blue is used for the output value “0” and the color red for the output value “1”.

#### Boolean Expression

Alternatively, a BF *f* can instead be represented as an algebraic expression (Fig. 1(c)) constructed with the *k* input variables which are combined via the logical operators AND (· or ∧), OR (+ or ∨) and NOT (′ or −). For example, the AND function and OR function of 2 input variables, *x*_1_ and *x*_2_, are given by the Boolean expressions, *x*_1_ · *x*_2_ and *x*_1_ + *x*_2_, respectively. Note that the “+” symbol does not correspond to working modulo 2, instead (1+1) has the value “1”, not “0”. In this work, the term *literal* refers to a Boolean variable (e.g., *x*_1_) or the complement of a Boolean variable (e.g., 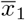). Throughout this work, *Boolean function* and *Boolean variables* refer to a *logical update rule* and to *inputs*, respectively, of nodes in a model.

#### Colored Boolean Hypercube

A visually illustrative representation of a BF is obtained by coloring a Boolean hypercube. A *k*-dimensional hypercube (*k*-cube) is composed of vertices and edges where each vertex is labelled by a string of *k* bits, and is connected to vertices with labels that differ from its label in exactly one bit. Two vertices connected by an edge are called “neighbors”. A *k*-cube thus has 2^*k*^ vertices, with each vertex having *k* neighbors. The total number of edges in a *k*-cube is (*k* × 2^*k*^)/2 (division by 2 removes the doubly counted edges). A BF may thus be represented by a *k*-cube in which each vertex is labelled by the input combination *x_k_ x*_*k*−1_*x*_*k*−2_…*x*_2_*x*_1_ (*x_i_* ∈ {0, 1}) and is colored with an output bit (0 or 1) (Fig. 1(d)).

#### Bias and Parity

In the Boolean modeling literature, the *bias p* of a BF refers to the *probability* for it to take the value 1 [2, 26]. Thus, taking all inputs as equally probable the bias *p* of a BF *f* with *k* inputs is the fraction of the 2^*k*^ rows of the truth table for which the output value of the BF is 1. In the computer science literature, the number of 1s in the output column of the truth table is called the *Hamming weight* (HW) of the BF. The HW is thus the *unnormalized bias P* of a BF. As a short hand, we will refer to the Hamming weight *P* simply as the *bias*. If a BF has odd or even bias *P*, the *parity* of the BF is said to be odd or even, respectively.

### C. Categorization of Boolean functions based on their bias and isometries

In his work, Feldman [28, 35] has classified BFs based on *D*, which in his notation is the number of inputs (or variables), and on *P* (or HW), the unnormalized bias. For notational consistency, we will use *k* instead of *D* hereafter and will denote the set of all BFs of given number of inputs *k* and bias *P* as *k*[*P*]. It is easy to see that the number of *k*[*P*] sets for a given *k* is 2^*k*^ + 1.

In addition, within any given *k*[*P*] set, Feldman [28, 35] introduced partitions using equivalence classes based on isomorphisms. Two BFs *f* and *g* are defined as *isomorphic* if they are identical up to permutations and negations of any of their input variables. In terms of the truth table, an isomorphism corresponds to permutations of columns associated with the inputs and to exchanging 0s and 1s within any columns. For example, the BF *f* = *x*_1_ ·(*x*_2_ + *x*_3_) is isomorphic to the BF 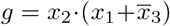. These transformations define equivalence classes of BFs within the *k*[*P*] set. For each class, we can choose the representative BF where the first occurrence of each variable arises both sequentially (with indices 1, 2, 3,…) and as a positive literal. Note also that every function in *k*[*P*] has a *complementary* function in *k*[2^*k*^ – *P*] which can be obtained via complementation of the corresponding Boolean expression (see e.g., [35]).

Visually, a BF in a *k*[*P*] set can be thought of as a *k*-cube wherein any *P* vertices are colored red (corresponding to output value 1) and the remaining 2^*k*^ – *P* vertices are colored blue (corresponding to output value 0). Note that on the Boolean hypercube, isomorphic elements of any arrangement of *P* red vertices (i.e., 1s) can be generated by rotation (i.e., permutation) and reflection (i.e., negation) about any axis (i.e., input variable) of the Boolean hypercube (i.e., function). For any given assignment of *P* “1”s on the *k*-cube, the total number of edges stemming from these *P* vertices is *kP*. Of these, some edges end at one of the other *P* – 1 vertices with value 1; we refer to this set of edges as *E*_11_. Similarly, we denote by *E*_01_ the remaining edges, ending at any of the 2^*k*^ – *P* other vertices having the value 0. We then have *E*_01_ + 2*E*_11_ = *kP* [36].

In sum, the introduction of the set *k*[*P*] and the classification scheme proposed by Feldman [28, 35] efficiently encapsulate the 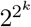 possible BFs with *k* inputs in terms of significantly fewer representative quantities and expressions. Interestingly, Reichhardt and Bassler [37], using concepts borrowed from isomer chemistry and group theory, have shown how to count the number of distinct representative non-isomorphic BFs in each *k*[*P*] set.

## III. COMPLEXITY MEASURES

### A. Minimal expressions and the Boolean Complexity

In the field of computer science, various propositions for measuring the *complexity* of BFs have been extensively studied [38–40]. In the present work, one definition we consider is due to Feldman [28]: it specifies the Boolean complexity of a BF *via* the complexity of the associated logical expression. In principle, there are an infinite number of logical expressions corresponding to a given BF. Feldman [28] focused on the shortest possible expression (in terms of the number of literals it is composed of), the so called *minimal formula* for a BF. Thereafter, Feldman defined the *Boolean complexity* of a BF to be the number of literals in its minimal formula [28, 38].

Note that Boolean expressions such as the canonical disjunctive normal form (DNF), the canonical conjunctive normal form (CNF), the minimal DNF and the minimal CNF, are widely-used to represent BFs. However, these forms of a BF need not provide the minimal formula in terms of Boolean complexity as defined by Feldman [28].

To illustrate the minimization of a Boolean expression, consider the following BF with 3 inputs:

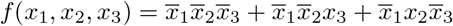

which contains 9 literals in the canonical DNF form. By applying the laws of Boolean algebra, we can simplify the expression as follows:

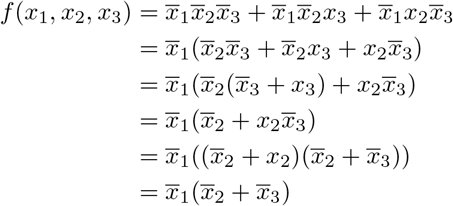

In the above simplification, we first factor out 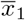 from the canonical DNF expression. Next, the expression within the braces is simplified further by factoring 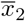. As 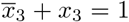, the above factorization leads to the simplified form 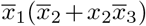. To further simplify the obtained expression within the braces, we next apply the distribution property over the OR (+) operator. As 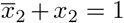, we finally obtain the expression 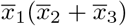 which is irreducible and hence is the minimal expression. This expression has a Boolean complexity equal to 3 as it has 3 literals. However, note that the minimal DNF for the above BF is 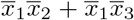, which has 4 literals, and factorization of this expression is necessary to obtain the minimal expression with 3 literals for the above BF.

#### Computing the Boolean complexity of a function

As shown above, the definition of Boolean complexity of a BF necessitates its reduction to a minimal (equivalent) formula. Obtaining a minimal formula for a given BF or expression is a computationally hard problem [41]. In practice, for more than a handful of variables, one has to resort to heuristic algorithms such as the QMV proposed by Vigo [42] for reducing expressions. Thus, barring exceptions, one can only obtain an upper bound on the Boolean complexity for BFs with several inputs. In this work, to obtain the factorized minimal expression of a BF, we employ the logic synthesis software “ABC” [43, 44]. To improve the estimated Boolean complexity of a BF, we give as input to the ABC software four types of Boolean expressions, namely the full DNF, the full CNF, the Quine-McCluskey minimized DNF expression [45, 46] and the Quine-McCluskey minimized CNF expression, corresponding to the same BF. As a result, 4 output Boolean expressions are obtained of which the one with the least number of literals is chosen as the minimal equivalent expression of the BF. The number of literals in this expression is then our estimate of the Boolean complexity of that BF.

### B. Average sensitivity of Boolean functions

For a BF *f* with *k* inputs, the *sensitivity S_f_* (**x**) for a given assignment of the input variables **x** = (*x*_1_ = *a*_1_, *x*_2_ = *a*_2_,…, *x_k_* = *a_k_*) is defined as the number of *neighbors* **y** of **x** for which the output *f*(**y**) is different from *f*(**x**) [26, 34]. Note that the assignments **y** and **x** are neighbors if their Hamming distance is 1, that is, they differ in exactly one of the *k* variables. Mathematically, the sensitivity is given by:

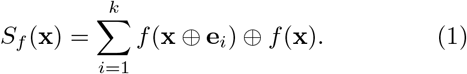

In the above equation, φ is the XOR operator and **e**_*i*_ ∈ {0,1}^*k*^ denotes the unit vector corresponding to having input variable *x_i_* = 1 and all others *x_i_* = 0. In terms of the hypercube representation, the sensitivity of any given vertex (a particular input vector) *i* on the hypercube is the number of its *k* neighbors that have output values which differ from the output value at vertex *i*. The total sensitivity of *f* is the sum of the sensitivities over all the vertices of its corresponding Boolean hypercube. Note that the total sensitivity of *f* is twice the number of edges whose two ends are vertices with complementary output values in the *k*-cube (i.e., 2*E*_01_). Notably, the sensitivity is also a measure of complexity of BFs [39].

It is easy to see that two BFs which are isomorphic to each other have the same average sensitivity. Since the operations of rotations or reflections about any of the axes of the hypercube do not change the number of “red” or “blue” neighbors of any vertex, the total sensitivity of the BF is preserved for all of its isomorphic forms. Moreover, a BF and its complement belonging to sets *k*[*P*] and *k*[2^*k*^ – *P*], respectively, also have the same average sensitivity. This is so because under complementation of the BF, the “red” and “blue” vertices of the *k*-cube are exchanged, thereby leaving the number of edges *E*_01_ in the *k*-cube unchanged.

Averaging the sensitivity, *S_f_*(**x**), over all possible input vectors or rows of the truth table gives the *average sensitivity* of a BF, that is:

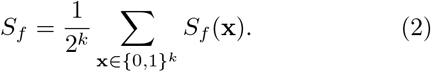

This is essentially the total sensitivity averaged over the 2^*k*^ vertices in the *k*-cube. The average sensitivity *S_f_* takes values in the range [0, *k*]. Further, it follows from the above definition that BF with lower average sensitivity are more *robust* to changes in their input variables [26].

## IV. BIOLOGICALLY MEANINGFUL TYPES OF BOOLEAN FUNCTIONS

In the literature, various types of BFs have been introduced of which some may be particularly suited to biological modeling [29]. In this section, we describe a number of types of *biologically meaningful* BFs and give some of their properties.

### A. Effective functions

A BF may possess inputs which are mute or *ineffective* in the following sense: altering the binary state of an ineffective input, while keeping the state of other inputs unchanged, never alters the output value of the BF. Biologically, a regulatory element can be considered to be an *effective* regulator of the expression of a gene if and only if there exists some input condition wherein the modulation of the regulator alters the expression of the gene. If such a condition does not exist, then that regulator (or input) can be considered to be *ineffective* as it plays no role in regulating the gene under consideration. It follows that all inputs (or regulators) of a biologically meaningful BF should be effective [29].

#### Definition 1

(Effective function (EF)). A BF *f* with *k* inputs is an EF if and only if:

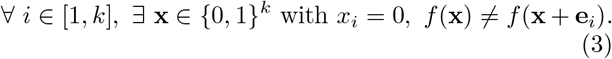

Here **e**_*i*_ ∈ { 0, 1 }^*k*^ denotes the unit vector associated to the component of index i. Simply put, all the inputs of an EF are effective.

#### Property IV.1.

EFs with *k* inputs have Boolean complexity ≥ *k*.

*Proof*: For a BF *f* to be effective, the *k* different input variables must appear at least once in the minimal expression or formula for the BF. This implies that the number of literals in the minimal expression or Boolean complexity is ≥ *k*.

### B. Unate functions

A regulatory element may act as an activator or an inhibitor of the expression of a target gene, and this information on the nature of the interaction between the regulator and its target gene can also be incorporated into the BF. The BFs which can account for this *sign* of regulatory interactions are known as Unate functions.

#### Definition 2

(Unate function (UF) [30]). A BF *f* with *k* inputs is said to be increasing monotone (activating) in an input *i* (or variable *x_i_*) if:

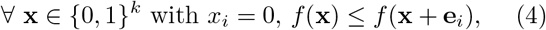

and decreasing monotone (inhibiting) in an input *i* (or variable *x_i_*) if:

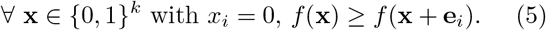

A BF *f* with *k* inputs is said to be a sign-definite or *unate function* (UF), if each input *i* = 1, 2,…, *k* is increasing monotone (activating) or decreasing monotone (inhibiting). Therefore, UFs with *k* inputs can be further classified into different combinations of activating and inhibiting inputs (Fig. S2). A BF *f* with *k* inputs is said to be a positive (respectively negative) UF, if every input *i* = 1, 2,…, *k* is increasing monotone (respectively decreasing monotone) [30, 47].

#### Property IV.2.

A UF can be represented by an expression in disjunctive normal form (DNF) in which all occurrences of any specific input variable (more precisely, literal) are either negated (i.e., negative input) or non-negated (i.e., positive input) [30, 47].

We next show that a UF whose input acts both as an activator and an inhibitor is ineffective in that input.

#### Property IV.3.

If an input *i* of a UF *u* acts as both an activator and an inhibitor, then input *i* is ineffective.

*Proof*: An input *i* can act as both an activator and an inhibitor, if and only if, the input *i* satisfies the equality condition in Eqs. 4 and 5, respectively, for all input vectors **x**. However, this is precisely equivalent to the condition for an input *i* to be ineffective.

In Appendix A and B, we present some properties associated with combining two independent BFs including UFs. Note that by ‘all sign combinations’ of UFs in any table or figure caption, we mean that every input is either activating or inhibiting. For instance, a function with *k* = 3 inputs could have 3 activators and no inhibitors, or 2 activators and 1 inhibitor, or 1 activator and 2 inhibitors, or no activators and 3 inhibitors (Fig. S2), and all such possibilities are considered for a UF of *k* inputs.

### C. Canalyzing functions

#### Definition 3

(Canalyzing function (CF) [2]). A BF *f* with *k* inputs is said to be canalyzing in an input *i* (or variable *x_i_*) if and only if:

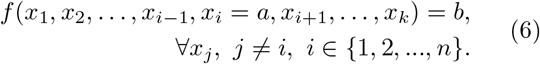

In the above equation, *a* and *b* can take values 0 or 1, *a* is the canalyzing input value and *b* is the canalyzed value for input *i*. A BF *f* is a *canalyzing function* (CF) if at least one of its *k* inputs satisfies the canalyzing property. Furthermore, for a CF, the number of distinct inputs that satisfy the canalyzing property is referred to as its *canalyzing depth*. For a CF with *k* inputs, the canalyzing depth can have an integer value in the range [1, *k*].

### D. Nested Canalyzing functions

#### Definition 4

(Nested Canalyzing function (NCF) [21, 32, 48, 49]). A BF *f* with *k* inputs is *nested canalyzing* with respect to a permutation *σ* on its inputs {1, 2,…, *k*} if:

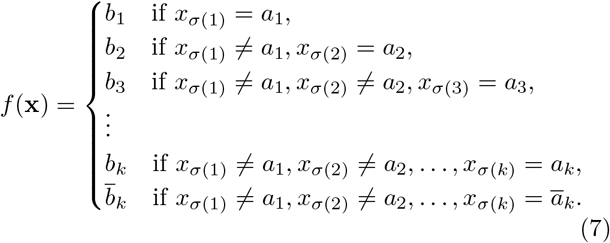

In the above equation, *a*_1_, *a*_2_,…, *a_k_* are the canalyzing input values and *b*_1_, *b*_2_,…, *b_k_* are the canalyzed output values for inputs *σ*(1), *σ*(2),…, *σ*(*k*) in the permutation *σ* of the *k* inputs. Here, *ā_k_* and 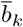 are the complements of the Boolean values *a_k_* and *b_k_*, respectively. We remark that Szallasi and Liang [48] had called these functions “hierarchically canalyzing” and subsequently, Kauffman [21] called them “nested canalyzing”.

#### Property IV.4.

A NCF with *k* inputs can be represented as a Boolean expression with exactly *k* literals as follows [49, 50]:

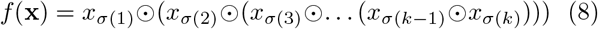

where *σ* is a permutation on the inputs {1, 2,…, *k*} and ⊙ represents either the AND (∧) or OR (∨) operator.

*Corollary*: The factorization of the DNF of NCFs reveals the *hierarchical* structure of the NCFs.

#### Property IV.5.

NCFs with *k* inputs have Boolean complexity equal to *k*.

*Proof*: In Eq. 8, each variable or literal in the expression for an NCF appears exactly once, thus the Boolean complexity of a NCF is equal to *k*. In Appendix B, we also show that NCFs are effective and unate functions. Thus, NCFs have the *minimum* Boolean complexity among EFs.

### E. Read-once functions

#### Definition 5

(Read-once function (RoF) [51]). A BF of *k* variables is a read-once function (RoF) if it can be represented by a Boolean expression, using the operations of conjunction, disjunction and negation, in which every variable appears exactly once [51]. RoFs have also been studied under the name of fanout-free functions in the computer science literature [52]. Mathematically, a BF *f* = *f*(*x*_1_, *x*_2_,…, *x_k_*) is a RoF if and only if there is a permutation *σ* on its inputs {1, 2,…, *k*} such that

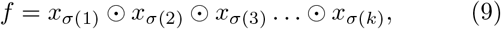

where ⊙ represents the AND (∧) or OR (∨) operator and all parentheses have been omitted in the expression. Note that there are no restrictions on the placement of those parentheses. For instance, the expressions for the RoFs, *x*_1_*x*_2_(*x*_3_ + *x*_4_) and *x*_1_(*x*_2_*x*_3_ + *x*_4_), have the same 4 variables but different placement of parentheses, while the expressions for the RoFs, *x*_1_*x*_2_(*x*_3_ + *x*_4_) and *x*_1_(*x*_2_ + *x*_3_ + *x*_4_), have the same 4 variables but different placement of parentheses and different positions of the AND or OR operators. Moreover, we show in Appendix B that the RoFs are effective and unate since each variable appears exactly once in the Boolean expression for a RoF. We next show that NCFs are a subset of the RoFs.

#### Property IV.6.

For any *k*, NCFs are a subset of RoFs.

*Proof*: Comparing the expression for NCFs (Eq. 8) with the expression for RoFs (Eq. 9), it is evident that NCFs form a subset of RoFs. Simply stated, all NCFs are RoFs but all RoFs are not NCFs. Henceforth, we refer to the subset of the RoFs which are not NCFs as the ‘non-NCF RoFs’. To the best of our knowledge, RoFs (excluding NCFs) have not been considered in the biological literature.

#### Property IV.7.

RoFs with *k* inputs have the minimum Boolean complexity *k* among all the EFs.

*Proof*: Since RoFs are constructed such that each variable or literal in the expression for a RoF (Eq. 9) appears exactly once, the Boolean complexity of a RoF is equal to *k*. Further, using property IV.1, it is clear that the RoFs satisfy the minimum equality condition, and hence, have the minimum Boolean complexity among all EFs. In sum, RoFs correspond exactly to the set of EFs with *minimum* Boolean complexity.

For the sake of compactness, we will represent RoFs which are equivalent up to isomorphisms (i.e., permutations of indices and complementation of variables) via a single representative BF or expression. Furthermore, we will classify RoFs into *k*[*P*] sets based on the number of inputs *k* and bias *P* [28]. In other words, we capture the complete set of RoFs in different *k*[*P*] sets via representative RoFs wherein each representative RoF captures all RoFs that are equivalent up to isomorphisms. For example, among BFs with *k* = 4 inputs, there are 10 representative RoFs up to isomorphisms which are:

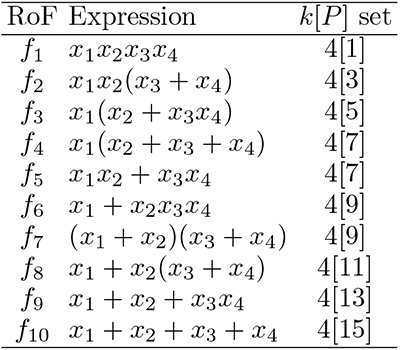

| RoF | Expression | $k[P]$ set |
| --- | --- | --- |
| $f_1$ | $x_1x_2x_3x_4$ | 4[1] |
| $f_2$ | $x_1x_2(x_3 + x_4)$ | 4[3] |
| $f_3$ | $x_1(x_2 + x_3x_4)$ | 4[5] |
| $f_4$ | $x_1(x_2 + x_3 + x_4)$ | 4[7] |
| $f_5$ | $x_1x_2 + x_3x_4$ | 4[7] |
| $f_6$ | $x_1 + x_2x_3x_4$ | 4[9] |
| $f_7$ | $(x_1 + x_2)(x_3 + x_4)$ | 4[9] |
| $f_8$ | $x_1 + x_2(x_3 + x_4)$ | 4[11] |
| $f_9$ | $x_1 + x_2 + x_3x_4$ | 4[13] |
| $f_{10}$ | $x_1 + x_2 + x_3 + x_4$ | 4[15] |

Among the above-mentioned 10 RoFs with *k* = 4, *f*_1_, *f*_2_, *f*_3_, *f*_4_, *f*_6_, *f*_8_, *f*_9_ and *f*_10_ are also NCFs.

#### Property IV.8

(Generation of representative RoFs). The following is a recursive scheme to generate all RoFs with *k* inputs, i.e., RoF(k), starting from RoFs with 1 input (*RoF*(1)). To do so, one can use the fact that the parentheses in the logical expression of a function in *RoF*(*k*) define two sub-parts separated by an AND or an OR operator. Such a decomposition splits the *k* variables into two sets, and thus, any function in *RoF*(*k*) can be decomposed into at least one of the following types:

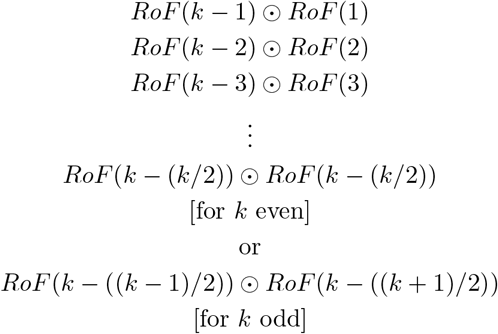

where ⊙ corresponds to the AND (∧) or OR (∨) operator. Such a decomposition allows one to enumerate all elements of *RoF*(*k*) recursively.

The above algorithm, does not exclusively return the representative RoFs, but also some of their permutations. To retain only the representative RoFs, we iteratively walk through each RoF in a *k*[*P*] set given by the above scheme, generate its list of permutations, and remove any RoFs in the *k*[*P*] set which belong to that list. We separately store the RoF with which we started out, essentially retaining just one of the permutations.

## V. OVERLAPS BETWEEN TYPES OF BOOLEAN FUNCTIONS AND IMPORTANT PROPERTIES

In the preceding section, we described multiple types of BFs that are expected to have biological relevance. Using exhaustive enumeration of BFs with *k* ≤ 4 inputs, we next explore quite systematically the relationships between the different types of biologically meaningful BFs. This computational exploration has further led us to conjecture some important properties for these different types of BFs, and subsequently, we have been able to provide analytical proofs. To the best of our knowledge, such a combined delineation of the different types of biologically meaningful BFs in the space of all 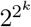 BFs has not been carried out previously.

By computational enumeration up to *k* ≤ 5, it is seen that the fraction of EFs within the space of all possible 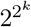 BFs with *k* inputs increases with increasing *k*. In contrast, the fraction of UFs and CFs decreases with increasing *k* (Tables S1 and S2, Figs. 2 and S1), and tend to 0. Table S3 also gives the fraction of EFs, UFs and CFs with even bias for *k* ≤ 5, and these fractions tend to 0.5 for increasing *k*. Table S4 also gives the number and fraction of ‘effective and unate functions’ (EUFs) with even bias for *k* ≤ 5 and different combinations of activators and inhibitors.

**FIG. 2.**
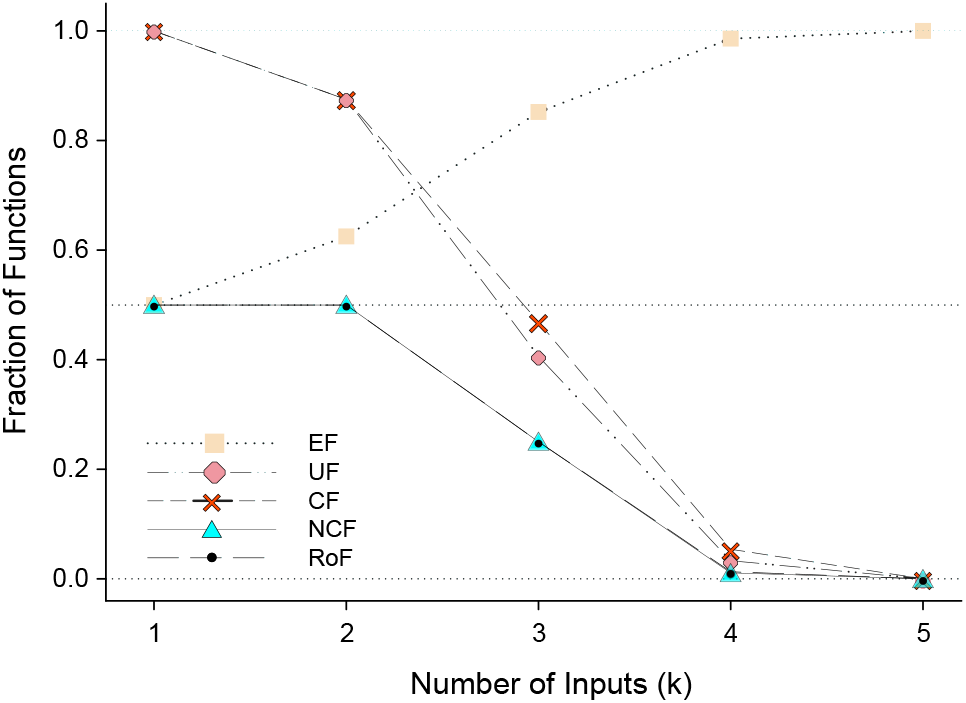
The fraction of biologically meaningful types of Boolean functions (BFs) among all BFs for a given number of inputs *k* ≤ 5. Here, EF corresponds to effective functions, UF to unate functions (all sign combinations), CF to canalyzing functions, NCF to nested canalyzing functions, and RoF to read-once functions.

Further, by computational enumeration up to *k* ≤ 10, it is seen that the fraction of RoFs, NCFs and non-NCF RoFs among all BFs with exactly *k* inputs decreases and tends to 0 with increasing *k* (Table S5, Fig. 2). Notably, we find that the fraction of NCFs among RoFs for a given value of *k*, decreases with increasing *k* (Table S5). In Fig. S3, we show the distribution of RoFs, NCFs and non-NCF RoFs with *k* = 4, 5, 6, 7 and 8 inputs, across different values of their bias *P*.

Fig. 3(a) serves as a visual guide to the overlaps between the different types of BFs. We begin by dividing the space of all BFs into 2 equal parts based on the parity of the bias. It is easy to see that the number of BFs with odd bias is equal to that with even bias. Next we observe that all ineffective BFs (which have at least one ineffective input, i.e., IEFs) lie in the even bias half. This raises the question as to whether all IEFs have even bias, and we theoretically prove that this is indeed the case below.

**FIG. 3.**
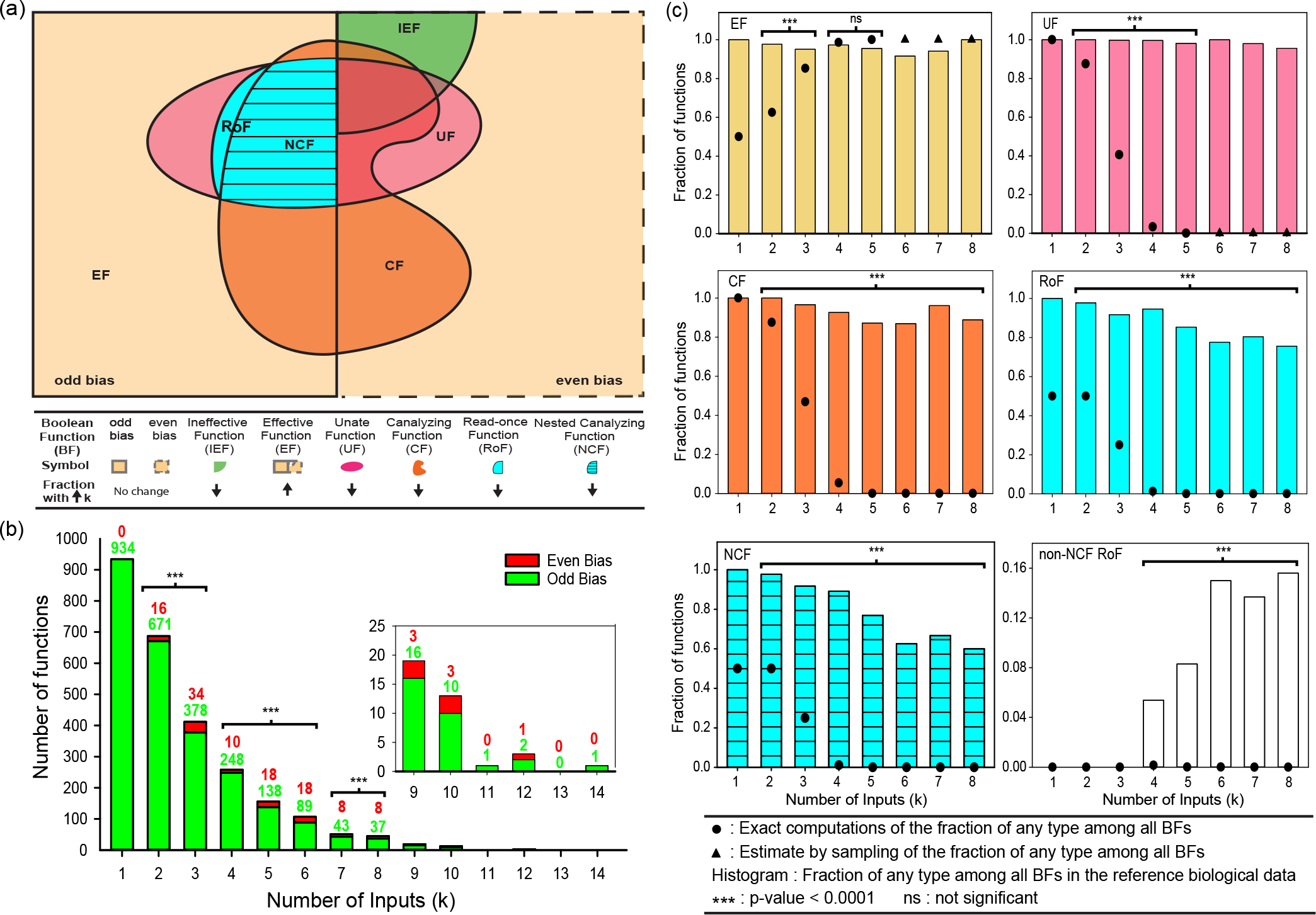
**(a)** The schematic figure depicts the overlap between different types of biologically meaningful Boolean functions (BFs) with *k* = 4 inputs. While this figure is not drawn to scale, the sizes of the sets corresponding to different types of BFs with 4 inputs and their intersections respect the order of the actual values. The legend gives the correspondence between shapes with specific color to different types of BFs. Arranging the different types of BFs with 4 inputs (which are *not* mutually exclusive) based on their sizes in a descending order gives: EF > Odd bias = Even bias > CF > UF > RoF > NCF. The up (or down) arrows in the legend depict the increase (or decrease) in the fraction of BFs that belong to a specific type as the the number of inputs *k* increases (see Table S2 for the exact numbers). **(b)** The in-degree distribution (or number of inputs) for nodes in the reference biological dataset of 2687 BFs from 88 models of living systems. The number of BFs decreases when the in-degree grows. Further, the number of BFs with odd bias (indicated in ‘green’) are significantly greater than those with even bias (indicated in ‘red’) in the dataset for *k* < 10. **(c)** The plots show that the biologically meaningful types, EF, UF, CF, RoF, NCF and non-NCF RoF, for a given number of inputs *k* ≤ 8 are over-represented in the reference dataset (except for EFs with *k* = 4, 5). For a given *k*, at least 60% of the reference biological BFs belong to the considered types (except for non-NCF RoF). Note that dot symbols which appear to coincide with the x-axis are very small non-zero numbers (except for non-NCF RoFs with *k* = 1, 2, 3). The raw data associated with these 6 plots along with results from the statistical test for over-representation are included in Tables S6, S7 and S8.

### Property V.1.

The bias *P* of a BF with *m* ineffective inputs is a multiple of 2^*m*^.

*Proof*: Consider the truth table of a BF *f* and assume the input variable *x_i_* is ineffective. Then each line with *x_i_* = 0 can be uniquely paired with the corresponding line having *x_i_* = 1 where all other variables are unchanged. Since the output is the same in both of these lines, one has either no 1s or two 1s in the output. Summing over all of the truth table (lines coming in pairs) will thus lead to an even number of 1s.

*Corollary*: It immediately follows that a BF with an odd bias is effective.

Moving on to the UFs, we find that they are rather evenly distributed across even and odd biases, and overlap with the IEF domain (Fig. 3(a)). Note that the space of UFs (for a given number of *k* inputs) allows for all possible numbers of activators and inhibitors (Fig. S2). We observe that not all UFs are EFs (see Property IV.3). The CFs, like the UFs, are rather equally shared across even and odd biases. CFs overlap with the IEFs, EFs and UFs (Fig. 3(a)).

We next examine the space of RoFs. Interestingly, for the BFs with *k* = 4 inputs, RoFs lie in the odd bias half (Fig. 3(a)). This warrants the conjecture that all RoFs have odd bias, and we show below that this is indeed the case.

### Property V.2.

RoFs have odd bias.

*Proof*: Consider the basic case of a RoF with 1 input. Clearly, the BFs in *RoF*(1) have output vector (0, 1) or (1, 0), both of which have odd bias. Next, let us hypothesize that BFs in *RoF*(*j*) ∀ *j* ∈ {1, *k*} have odd bias. We now refer to Property A.3 in Appendix whereby the combination of two BFs with odd bias results in a BF with odd bias. Next by induction, the RoF with *k* + 1 inputs is given by, 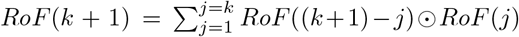 by Property IV.8. Since the two functions *RoF*((*k* + 1) – *j*) and *RoF*(*j*) have odd bias, using Property A.3, the function *RoF*(*k* + 1) will also have an odd bias.

Moving to the NCFs, we see in Fig. 3(a) that NCFs lie within the space of RoFs, and this is actually true for any number of inputs *k* (see Property IV.6). It turns out that NCFs are a strict subset of RoFs when *k* ≥ 4, or in other words, there are non-NCF RoFs for *k* ≥ 4. Also note that NCFs have odd bias as they are a subset of RoFs. For an alternate proof, see Property B.3 in Appendix.

## VI. ENRICHMENTS IN THE BIOLOGICAL DATA

In this section, we report our observations on the relative abundance and associated statistical significance of the different types of BFs in a compiled dataset of BFs from reconstructed models. In Appendix C, we provide details on the compilation and curation of this reference biological dataset of 2687 BFs from 88 published models. The in-degree distribution for these 2687 BFs comprising the dataset analyzed here is shown in Fig. 3(b). As the number of inputs *k* to a BF increases, the number of nodes or functions with *k* inputs decreases very rapidly (Fig. 3(b)).

The key methodology hereafter consists in focusing on the relative abundances of the different types of BFs (and the statistical significance thereof) when comparing the ensemble of all BFs to the ensemble composed of our reference dataset. An enrichment, if it is statistically significant, is suggestive of some level of selection pressure on the BFs present in the biological networks. For details about the statistical tests, see Appendix D.

### A. Enrichment in types when comparing to the ensemble of random Boolean functions

Fig. 3(b) shows the abundances of the BFs as a function of in-degree in the reference biological dataset of 2687 functions. From Fig. 3(b), we observe that for each input 1 ≤ *k* ≤ 8, the odd bias BFs are dominant and statistically enriched in the reference dataset. It is not immediately apparent why BFs with odd bias should be preferred over BFs with even bias as in theory biologically meaningful BFs with even bias also exist (see Fig. 3(a)).

Fig. 3(c) shows the relative abundances in that dataset for different BF types, namely EFs, UFs, CFs, RoFs, NCFs, and non-NCF RoFs. Also shown in the figure are the values of these relative abundances in the ensemble of random BFs.

We have performed statistical tests to determine whether the relative abundances in the reference biological dataset are larger than expected (one-sided p-values) under the null hypothesis that the reference BFs are drawn from the ensemble of random BFs. We find that most of these types are indeed more abundant than expected (see stars above the bars in Fig. 3(c)). The main exception is the EF type of function; that feature can be understood by the fact that random functions are typically effective (see Fig. 2). For the detailed p-values, refer to Table S8. From the enrichment ratios shown in Table 1, it is clear that the RoF, NCF and also the non-NCF RoF types are all strongly enriched.

**TABLE 1.**
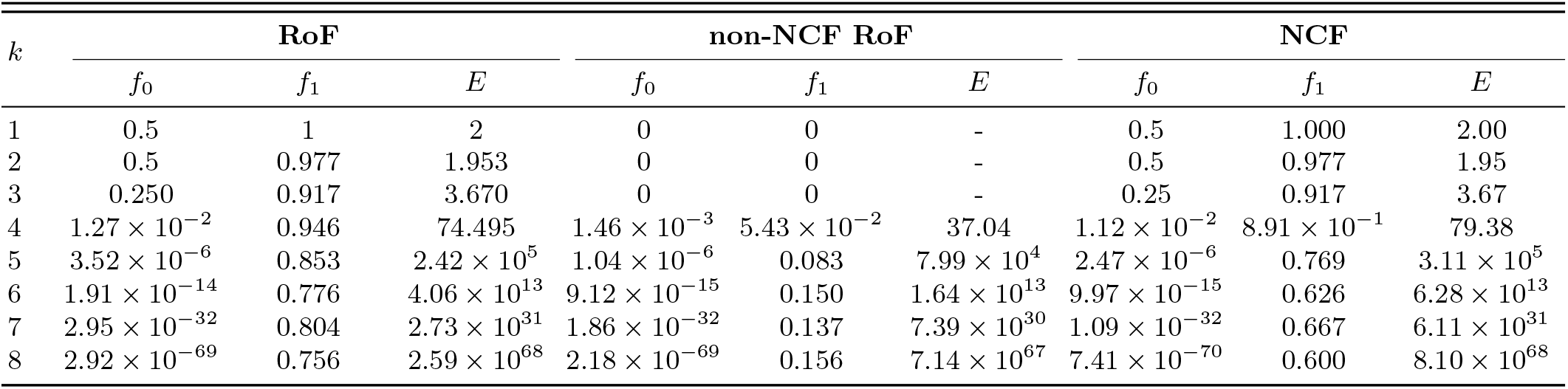
Fractions of functions that are RoFs, non-NCF RoFs or NCFs, in the space of all 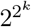 BFs (*f*_0_) or in the reference biological dataset (*f*_1_). *E*(= *f*_1_/*f*_0_) is the enrichment ratio; it indicates the extent of the over-representation of such functions in the reference dataset. Over-representation is highest for NCFs but clearly non-NCF RoFs are also highly over-represented. Computations are reported for functions with *k* ≤ 8 inputs.

### B. Relative enrichment in sub-types when comparing to random Boolean functions

Comparing the enrichments of the different types of biologically meaningful BFs can provide signatures of causes of enrichment. For instance, if selection operated only in favor of unateness, each sub-type therein (NCF, RoF or non-NCF RoF) would be expected to have a relative abundance (proportion within UF) that is the same whether one considers the reference biological dataset or the ensemble of random BFs. In effect, the proportions of different sub-types of BFs in the two ensembles points to which factors drive the different enrichments. We thus developed a way to test the hypothesis that a sub-type enrichment is solely due to the enrichment in one of its englobing types.

First, we consider the enrichment ratios of NCFs and RoFs in the three types of BFs: odd bias, EFs and UFs. From the Table S9, it is clear that for *k* > 2, the relative enrichment ratios (when comparing the observed to the expected under the null hypothesis) of both the NCFs and RoFs are much greater than 1, implying that the enrichment of these sub-types does not follow from the enrichment of their super-sets: biological selection *solely* in favor of being odd biased, effective or unate is not consistent with the enrichments found of the NCFs or RoFs in the reference dataset, some other factors are also at work.

Since NCFs are a subset of CFs, we can also ask whether canalyzation is the factor driving the enrichment of NCFs. Since the relative enrichment ratios are high and the p-values low (see Table S10), we can conclude that selection for canalyzation alone does not explain the enrichment observed for NCFs. Furthermore, we can ask whether it is minimum Boolean complexity, i.e., the fact that a function is a RoF, that drives the enrichment of NCFs (a sub-type of RoF). As shown in Table S10, the relative enrichment of NCF within RoF is quite modest, almost all *k* having *E_R_* values belonging to the range 1 to 2. Nevertheless our statistical method shows that these values are not consistent with 1 (absence of any enrichment) as shown by the p-values in Table S10, indicating that there must be some further cause of the enrichment of NCFs than that of belonging to RoF. We will examine the conjecture that a driving factor is our other measure of complexity, the average sensitivity (which is not constant within RoF).

We remark that the abundance of CFs and NCFs in reconstructed Boolean models of biological networks has been reported previously [2, 21, 22, 31, 50, 53–55]. Nevertheless, those studies did not provide statistical tests nor did they assess the relative enrichment in sub-types, a key methodology that provides insights into the factors likely to drive such enrichments.

## VII. ENRICHED FUNCTIONS IN BIOLOGICAL DATA HAVE MINIMUM COMPLEXITY

A plausible explanation for the enrichment of the RoFs and NCFs in the dataset is the *low complexity* of these functions. We have examined the different types of BFs using two notions of complexity (see section III). In terms of the first notion (Boolean complexity), the RoFs, of which NCFs are a subset, have in fact the minimum Boolean complexity among all EFs (see Property IV.7). RoFs and NCFs have the same Boolean complexity but interestingly they differ for the second measure of com-plexity, namely average sensitivity. Therefore the relative enrichment of NCFs and relative depletion of non-NCF RoFs within RoFs in the reference dataset can be possibly explained by using two measures of complexities which have relevance to regulatory logic in biological systems. This section thus examines more closely the properties of these two complexity measures. We also exploit the fact that for any bias, the minimum average sensitivity is obtained for a particular geometry of the “on” vertices of the k-dimensional hypercube (see section VII C). We will show that when the bias is odd this geometry corresponds to an NCF while if it is even the function is not effective.

### A. Correlation between Boolean Complexity and Average Sensitivity

To the best of our knowledge, the relationship between the two measures of complexity has not been explored. Fig. 4 portrays the relation between the Boolean complexity and average sensitivity for all the representative BFs with *k* = 4 inputs. As mentioned in section III A, obtaining the Boolean complexity is a computationally hard problem. We use the procedure outlined in section III A to compute the Boolean complexity of functions. The average sensitivity, unlike the Boolean complexity, can be computed exactly for a BF using Eq. 2. We perform a bivariate analysis of these two measures of complexity by computing their Pearson correlation coefficient (*ρ*) for all BFs with *k* = 4 inputs. The correlation between these measures turned out to be 0.812 (Fig. S4(d)), indicating a strong positive linear relationship between them. Looking closely at functions in the neighborhood of the ‘brown’ line (which highlights the minimum Boolean complexity 4 for EFs) in Fig. 4, we observe that:

- All EFs along this brown line have odd bias and are NCFs or non-NCF RoFs.
- At the bias *P* = 7, NCFs have a lower average sensitivity than the non-NCF RoFs, see Fig. S4(c) for a 2D projection.
- For each value of even bias, an IEF (Boolean complexity < *k*) has the minimum average sensitivity, see Fig. S4(c) for a 2D projection.

**FIG. 4.**
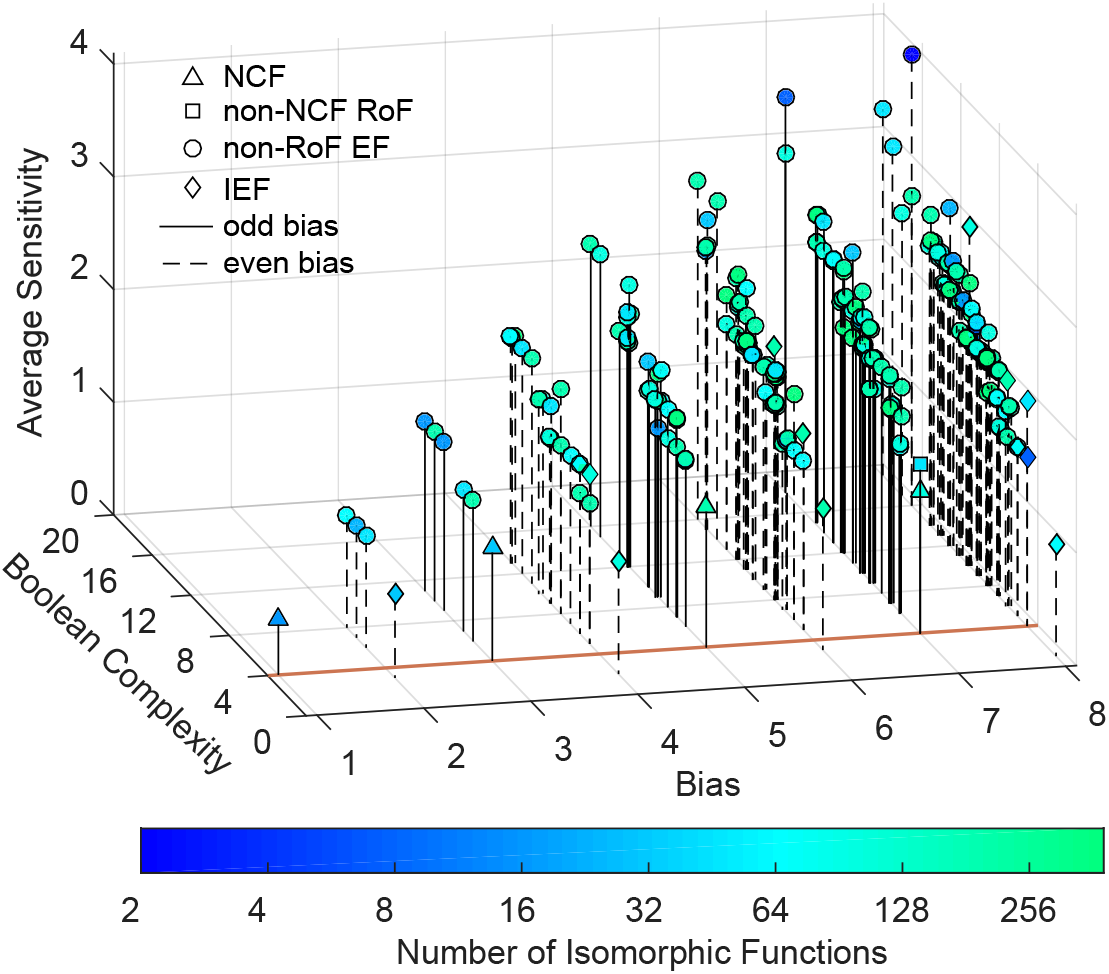
This figure depicts the variation of the average sensitivity and the Boolean complexity of Boolean functions (BFs) with *k* = 4 inputs as a function of bias *P*. Each point (with symbols circle, triangle, square or diamond) on the plot corresponds to one representative BF. The slight ‘jiggle’ at some points is added to resolve the overlapping representative BFs. Since these complexity measures are invariant under complementation of the BF, the bias values have been shown up to *P* = 8. The brown line drawn at the Boolean complexity 4 highlights the functions that possess the minimum Boolean complexity and are effective as well. Note that the NCFs are the only type of BFs that minimize both of the complexity measures. The high Pearson correlation coefficient (*ρ* = 0.812) between the average sensitivity and Boolean complexity indicates a strong correlation between these distinct measures of complexity. The two-dimensional (2D) projections of this three-dimensional (3D) plot are shown in Fig. S4.

These computational observations led us to two conjectures:

- When *P* is odd, NCFs have the minimum average sensitivity in a *k*[*P*] set.
- When *P* is even, the functions with minimum average sensitivity are ineffective with Boolean complexity < *k*.

We will prove these conjectures in the succeeding subsections.

### B. Maximizing the edges between *P* vertices of a *k*-cube minimizes the average sensitivity

Consider the problem of finding BFs which possess the lowest average sensitivity in a given *k*[*P*] set. In order to solve this problem, we will minimize the total sensitivity of the BF which is just a constant (2^*k*^) times larger. As mentioned in section III B, the total sensitivity on the hypercube is equal to twice the number of edges whose two ends are vertices with complementary bits. Recall the expression *kP* = *E*_01_ + 2*E*_11_ (see section IIB), where *E*_01_ is the total number of edges whose ends (vertices) have different values (colors), and *E*_11_ is the number of edges between the *P* vertices assigned the value “1”. Thus, the total sensitivity of a function in the set *k*[*P*] is 2*E*_01_. In order to minimize *E*_01_, we need to maximize *E*_11_ since *kP* is given. Note that the expression *kP* = *E***01** + 2*E*_11_ also implies that *E*_01_ + 2*E*_00_ = *k*(2^*k*^ – *P*). Thus, maximizing *E*_11_ will also maximize *E*_00_. This implies that finding a BF with minimum total sensitivity in *k*[*P*] set is equivalent to finding a BF that has the maximum number of edges among *P* vertices on the *k*-cube having the value 1 (the remaining 2^*k*^ – *P* vertices having the value 0).

### C. Edge-maximizing arrangement among *P* vertices: Defining “good sets”

In the work of Hart [36], the problem of finding a set of vertices on the hypercube that maximizes the number of edges between those same vertices has been solved. In fact this problem has been solved by multiple authors [56, 57] in different contexts in computer science. We chose to use Hart’s ideas due to it’s mathematical lucidity and easy visualization. Hart [36] had shown that there exists a class of vertices which solve the above mentioned problem, and he had referred to the class as a “good set” of vertices. A good set of *P* vertices on a *k*-cube where *P* < 2^*k*^, is defined recursively as follows:

i. If *P* = 1, we always have a good set.
ii. Otherwise, find *r* such that 2^*r*^ < *P* ≤ 2^*r*+1^. Select any (*r* + 1)-cube embedded in the *k*-cube. Then, select two *r*-cubes which are vertex disjoint subsets of the (*r* + 1)-cube. To select the *P* vertices, include first 2^*r*^ vertices by taking one of the *r*-cubes and include the remaining *P*– 2^*r*^ vertices by imposing that they form a “good set” containing *P* – 2^*r*^ vertices on the other r-cube.

By expressing *P* as a sum of powers of 2, i.e., 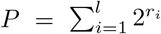, the resulting set of strictly increasing exponents {*r*_1_, *r*_2_,…, *r_l_*} gives the dimensions of the successive cubes to be used to define the good set.

The BFs in the set *k*[*P*] having minimum total sensitivity are those whose *P* vertices (with output value 1) and consequently the remaining 2^*k*^ – *P* vertices (with output value 0) both constitute “good sets”.

### D. A “good set” of odd vertices is a NCF

#### Conjecture

If a NCF in a *k*[*P*] set is considered in its hypercube representation, then the *P* vertices with output value 1 form a “good set”.

*Proof*: To see this, consider the logical expression of an NCF (Eq. 8) in a *k*[*P*] set. The *i^th^* canalyzing variable *x*_*σ*(*i*)_ determines which partition (of the possible *k* –(*i*–1) partitions, *i* – 1 variables having already been fixed) of a (*k* – (*i* – 1))-cube into 2 vertex disjoint (*k* – *i*)-cubes is to be canalyzed. Furthermore, the canalyzing input value *a_i_* (*x*_*σ*(*i*)_ = *a_i_*), fixes the outputs of the vertices of one of the two vertex disjoint (*k* – *i*)-cubes to the value *b_i_*. Repeating the above procedure recursively over *i* ∈ [1, *k*] gives the arrangement of 1s and 0s for an NCF on a *k*-cube. To obtain a NCF with a certain bias *P*, the *i*’s for which *b_i_* = 1 have to be chosen appropriately so that 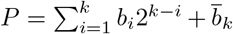, the last term arising because *P* is odd for NCFs.

The above procedure of setting the output values of *P* vertices to 1s and 2^*k*^ – *P* vertices to 0s on the *k*-cube is equivalent to obtaining a good set of *P* vertices, setting their output values to 1 and then setting the output of the remaining 2^*k*^ – *P* vertices to 0. This is true because:

i. the dimensions of the cubes whose vertices are to have the output value 1 are the same in either case (i.e., the set of exponents obtained by expressing *P* as a sum of powers of 2 is unique for a given *P*).
ii. when some *i*-cube is chosen to place the 1s, there is only one other *i*-cube, which (along with the chosen *i*-cube) constitutes 2 vertex disjoint subsets of a (*i* + 1)-cube. In both cases, this is an *i*-cube where the next set of 1s are placed.

This proves that the *P* vertices with output value 1 in a NCF constitute a good set. Therefore the NCFs have the minimum average sensitivity among all BFs in the set *k*[*P*] when the bias *P* is odd. Using two examples, we provide a visual proof of the above argument. (see Figs. 5 and S5)

**FIG. 5.**
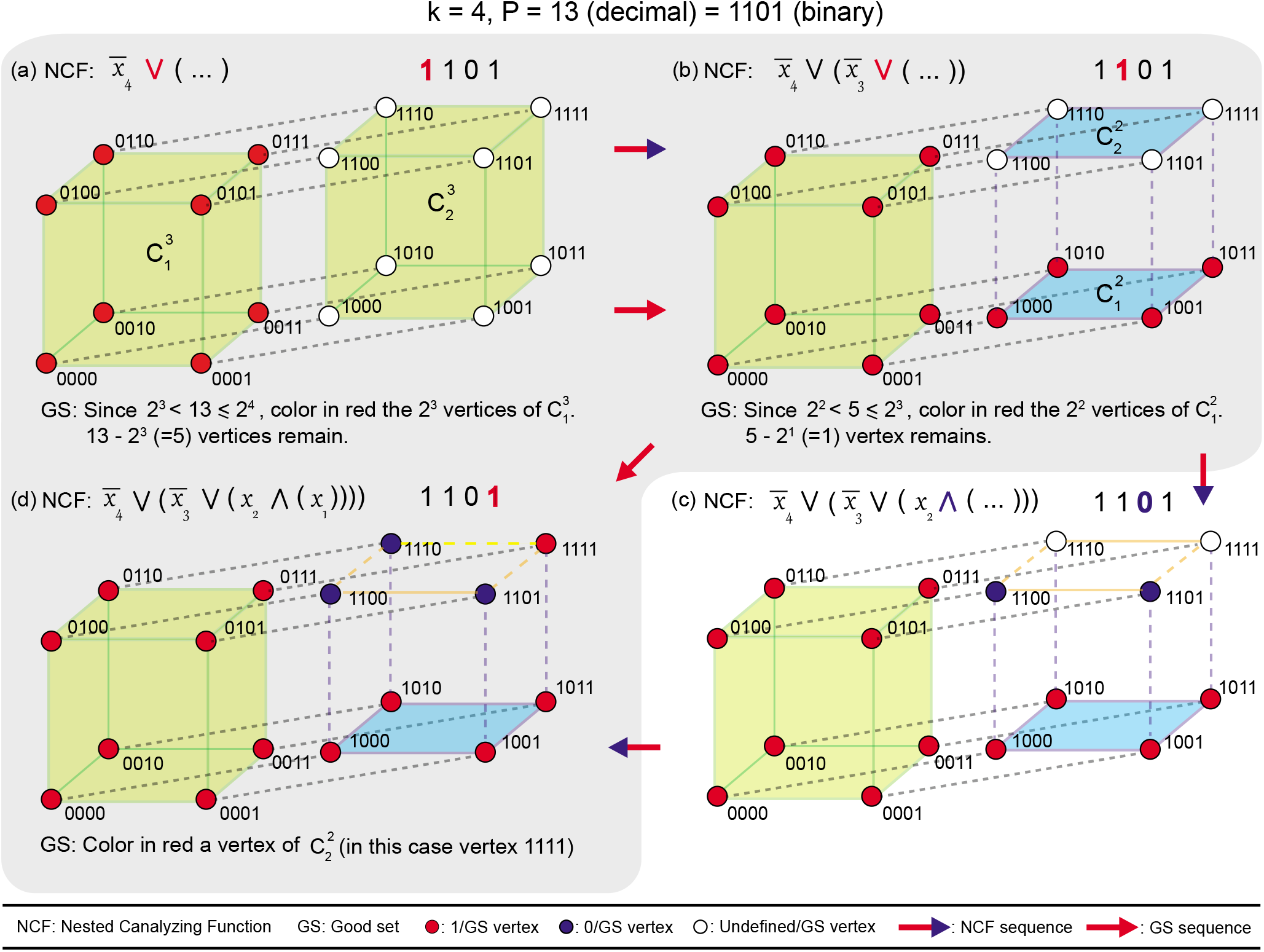
‘Good set’ (GS) for *P* vertices where *P* is odd on a *k*-dimensional hypercube is equivalent to a Nested Canalyzing Function (NCF) in that *k*[*P*] set. In parts (a), (b) and (d) shaded in grey, we show the recursive construction of a GS for *P* = 13 vertices in a 4-dimensional hypercube by coloring it’s vertices red, and in parts (a), (b), (c) and (d), we show the equivalence of that GS with 13 vertices to a NCF with bias 13. The vertices of the hypercube are labeled in the order *x*_4_, *x*_3_, *x*_2_, *x*_1_ wherein *x_i_* is 0 or 1. Here, 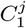 and 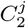 denote the two vertex disjoint *j*-dimensional hypercubes of the (*j* + 1)-dimensional hypercube. The ‘active’ bit in each part (a), (b), (c) and (d) is the colored bit in the binary representation of 13 in that part. **(a)** Since *P* = 13 lies between 2^3^ and 2^4^, 2^3^ vertices of either 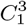 or 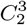 (here, 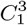) form part of the GS. This leaves 13 – 8 = 5 vertices to be colored red to complete the GS. This choice of 8 vertices in 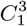 for the GS leads to the canalyzation of vertices labelled *x*_4_ = 0 to the output value 1. In this step, the active bit is 1 and as a result the ∨ operator follows the literal 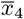. **(b)** Following the same procedure as in (a) for coloring the remaining 5 vertices of the GS leads to the choice of 4 vertices in 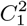. This leaves 1 vertex to be colored (which is the base case of the recursion to construct the GS). This choice of 4 vertices for the GS leads to the canalyzation of vertices with *x*_4_ = 1 and *x*_3_ = 0 to the output value 1. The active bit in this step is 1 and as a result the ∨ operator follows the literal 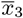. **(c)** For the corresponding NCF, the vertices with *x*_4_ = 1, *x*_3_ = 1 and *x*_2_ = 0 are canalyzed to the output value 0. The active bit in this step is 0 and as a result the ∧ operator follows the literal *x*_2_. **(d)** For the last step, any vertex in 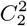 can be colored to complete the 13 vertices in GS, and we color here the vertex 1111. The vertex with *x*_4_ = 1, *x*_3_ = 1, *x*_2_ = 1 and *x*_1_ = 1 is canalyzed to the output value 1, and the remaining vertex is set to output value 0.

### E. A “good set” of even vertices has Boolean complexity strictly less than *k*

#### Conjecture

If an even number of vertices *P* having the output value 1 in the hypercube representation of a BF (in a *k*[*P*] set) forms a “good set”, then its Boolean complexity is strictly less than *k*.

*Proof*: The arrangement of *P* 1s and 2^*k*^ – *P* 0s on the *k*-cube in the case where *P* is even is almost the same as in a NCF, with the exception that the output values of the vertices of the last 1-cube (composed of 2 vertex disjoint sets of 0-cubes) will have the same output values *b_k_*. By direct computation, we have 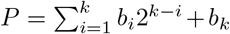 which is always even.

Now consider how to construct the DNF of a Boolean function (with bias *P*) defined by such a good set. Suppose that in the recursive construction of the good set one begins by assigning 1s to the vertices of a *j*-cube (*j* < *k*). The first clause of the DNF is then just the AND (product) of all the *k* – *j* literals involved to fill the vertices of that *j*-cube. If the next step of the recursive construction of the good set consists in assigning 1s to a *i*-cube (*i* < *j*), the second clause of the DNF will be the product of all *k* – *i* previous literals. We can thus iteratively construct the DNF for the BF represented by the given good set.

When output values of the vertices of the last 1-cube are to be fixed, both vertices have to be set to the same output value if *P* is even. Thus they will either contribute a term with *k* – 1 variables (if the output values are set to 1) or they will not contribute any term (if the output values are set to 0). Importantly, the variable which is missing in this clause is not present in any of the other clauses, therefore making that Boolean function ineffective in that input. In constructing such a function, there will be at most *k* – 1 variables in the Boolean expression. This implies that the resulting function has a Boolean complexity strictly less than *k*. See Fig. S6 for a visual proof of the above argument.

## VIII. DISCUSSION AND CONCLUSIONS

Ever since Kauffman [10, 11] proposed the Boolean modeling of gene regulatory networks, there has been considerable interest in this framework for studying the dynamics of biological networks. Subsequently, Kauffman [2] and others surmised that the logical rules or BFs determining the dynamics in such models of living systems are likely to have specific properties of biological relevance. Over the years, different types of “biologically meaningful” BFs have been introduced in the literature including effective (EFs) [29], unate (UFs) [30], canalyzing (CFs) [2] and nested canalyzing (NCFs) [21]. In particular, we introduced for the first time in a biological setting, the read-once functions (RoFs) [51] from the computer science literature. In this study, we have systematically investigated the occurrences and relationships among these different types of BFs in: (a) the space of all 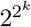 BFs, and (b) a reference dataset of 2687 BFs compiled from published Boolean networks [25, 58, 59] of biological systems.

Firstly, these biologically meaningful types of BFs represent a tiny fraction of the space of all BFs (see Fig. 2), and yet we find that they cover nearly all BFs found in our reference biological dataset (Fig. 3). In particular, we find that the odd bias BFs, UFs, CFs, RoFs and NCFs are highly over-represented in the reference biological dataset. Secondly, by studying the corresponding enrichments, we concluded that the enrichment of RoFs and NCFs in the reference dataset cannot be due only to odd bias, effectiveness or unateness. We remark that while the abundance of CFs and NCFs in biological networks has been previously reported in several publications [2, 21, 22, 31, 50, 53–55], we believe this is the first systematic study of 7 different types of BFs in a large curated reference biological dataset. Further, previous studies neither carried out statistical tests nor assessed the relative enrichments in sub-types, e.g. NCFs within CFs or RoFs, and in this respect, our study is able to shed light on possible driving factors.

Subsequently, we systematically examined two measures of complexity namely, Boolean complexity [28, 38] and average sensitivity [26, 34, 39], and found that their minimization largely accounted for the enrichments of RoFs and NCFs in the reference dataset. Inspired by the work of Feldman [28], we classified BFs into their respective *k*[*P*] sets, and thereafter examined abundances as a function of the three quantities: bias *P*, Boolean complexity and average sensitivity (Fig. 4). Based on this investigation, we find that RoFs minimize the Boolean complexity among EFs while NCFs minimize both the Boolean complexity and the average sensitivity for a given odd bias *P*. Though previous studies had noted that NCFs for a given number of inputs *k* have low average sensitivity [27, 49, 60] compared to other types of BFs, the *k*[*P*] classification (Fig. S4(c)) along with the invariance of the two complexity measures under complementation, led us to conjecture that NCFs achieve the *theoretical minimum* of this complexity measure in their *k*[*P*] set.

We then provided an *analytical proof* for this property of NCFs in *k*[*P*] sets by mapping the problem of minimizing the average sensitivity of a BF into one of edge minimization between two disjoint sets of vertices on the Boolean hypercube. Specifically, we leverage an important result by Hart [36] by formulating the problem for a given number of inputs *k* and fixed bias *P*. We showed that when *P* is odd, NCFs attain the absolute minimum of the average sensitivity over all possible BFs. When instead *P* is even, the minimum corresponds to BF that are not effective, though once the ineffective inputs are removed one recovers again precisely NCFs with fewer variables. As a result, NCFs have the surprising property that they minimize both Boolean complexity and average sensitivity.

The framework we used both supports and formalizes Kauffman’s [2] qualitative view in which “simplicity” should be a driver of the regulatory logic in biological systems. Kauffman argued that canalyzing functions were simpler than random functions, and therefore should be expected to arise quite frequently in biological systems [2, 53]. Our use of an extensive curated dataset generated from published Boolean models of biological networks enabled us to compare different notions of simplicity, and thereby confront Kauffman’s view to real data in a well defined quantitative framework. By identifying “simplicity” with minimum complexity defined in terms of either Boolean complexity or sensitivity of BFs, NCFs are the most simple. We can thus justify the much stronger preponderance of the NCF type in comparison to the CF type conjectured by Kauffman. We also note that sensitivity of BFs is directly related to their robustness to noise [6]. With that correspondence, we can conclude that NCFs for a given number of inputs *k* and given bias *P* have the *theoretically maximum robustness* to noise in the inputs. It is a posteriori natural to expect that average sensitivity as a measure of both complexity and robustness will be particularly relevant to Boolean models of gene regulatory networks.

Our methods and results have implications for the problem of model selection within the Boolean framework [61, 62]. By model selection we mean the process of selecting Boolean models from the ensemble of Boolean models which satisfy given constraints such as having specified steady states. During model selection, the preferential use of NCFs or RoFs could serve as a relevant criterion to constrain network reconstruction [61, 63]. In conclusion, minimum complexity functions are abundant and overwhelmingly dominate the regulatory logic in Boolean models of living systems.

## Appendix A: Combining two independent Boolean functions

Consider two *independent* BFs *f*_1_ and *f*_2_ with *k*_1_ and *k*_2_ inputs and bias *P*_1_ and *P*_2_, respectively. Here, the two BFs are independent in the sense that they have no input variables in common. The truth tables for the BFs *f*_1_ and *f*_2_ have 2^*k*_1_^ and 2^*k*_2_^ rows, respectively. A simple way to combine the two BFs is via AND or OR logical operators. We use the notation *f* = *f*_1_ ⊙ *f*_2_ where ⊙ is either the AND (∧) or OR (∨) operator. The procedure to generate *f* with 2^*k*_1_+*k*_2_^ rows in its truth table, by combining *f*_1_ and *f***2**, can be expressed compactly as follows:

#### Algorithm 1

Algorithm to combine two independent BFs *f*_1_ and *f*_2_

~~~
1: *f*[*r*] = 0, *r* ∈ [1, 2^*k*_1_+*k*_2_^]
2: *r* ← 1
3: **for** *i* ← 1 to 2^*k*_1_^ **do**
4:    **for** *j* ← 1 to 2^*k*_2_^ **do**
5:      *f*[*r*] = *f*_1_[*i*] ⊙ *f*_2_[*j*]
6:      *r* ← *r* + 1
7:    **end for**
8: **end for**
~~~

In Line 1 of the above algorithm, we initialize a vector *f* with 2^*k*_1_+*k*_2_^ elements to store the output values in each row of the truth table for *f* = *f*_1_ ⊙ *f*_2_. In Line 5 of the algorithm, the output value for the *i*^th^ row of *f*_1_ is combined with that of the *j*^th^ row of *f*_2_ to give the output value for the *r*^th^ row of *f*. For example, if BFs *f*_1_ and *f*_2_ with 1 input and 2 inputs, respectively, have output vectors (1,0) and (1,1,1, 0), respectively, then the output vector for the combined BF *f* = *f*_1_ ∧ *f*_2_ with 3 inputs is (1,1,1, 0, 0,0,0,0).

### Property A.1.

Let *f*_1_ and *f*_2_ have bias *P*_1_ and *P*_2_. Then *f*_1_ ∧ *f*_2_ (hereafter denoted as *f*_AND_) has bias equal to *P*_1_*P*_2_.

*Proof*: For every occurrence of 0 in the output vector of *f*_1_, the output vector of *f*_AND_ will also be 0. For every occurrence of 1 in the output vector of *f*_1_, there will be *P*_2_ occurrences of 1 in the output vector of *f*_AND_. Thus, for *P*_1_ occurrences of 1 in the output vector of *f*_1_, there will be *P*_1_*P*_2_ occurrences of 1 in the output vector of *f*_AND_.

### Property A.2.

Let us denote *f*_1_ ∨ *f*_2_ by *f*_OR_, then *f*_OR_ has bias equal to 2^*k*_1_^ *P*_2_ + 2^*k*_2_^*P*_1_ – *P*_1_*P*_2_.

*Proof*: For every occurrence of 0 in the output vector of *f*_1_, there will be *P*_2_ occurrences of 1 in the output vector of *f*_OR_. Thus, the contribution to the 1’s in the output vector of *f*_OR_ from 0’s in the output vector of *f*_1_ is (2^*k*_1_^ – *P*_1_)*P*_2_. For every occurrence of 1 in the output vector of *f*_1_, there will be 2^*k*_2_^ occurrences of 1’s in the output vector of *f*_OR_. Thus, the contribution to the 1’s in the output vector of *f*_OR_ from 1’s in the output vector of *f*_1_ is 2^*k*_2_^ *P*_1_. In total, the number of 1’s in the output vector of *f*_OR_ is equal to (2^*k*_1_^ – *P*_1_)*P*_2_ + 2^*k*_2_^ *P*_1_ = 2^*k*_1_^ *P*_2_ + 2^*k*_2_^ *P*_1_ – *P*_1_*P*_2_.

### Property A.3.

Given the bias parities (even or odd) of *f*_1_ and *f*_2_, the two previous results show that both *f*_AND_ and *f*_OR_ have *odd* parity (i.e., their bias is odd), if and only if *P*_1_ and *P*_2_ are both odd.

## Appendix B: Additional properties of biologically meaningful types of Boolean functions

Here, we present some additional properties for UFs, NCFs and RoFs.

### Unate functions

#### Property B.1.

If *u*_1_ and *u*_2_ are UFs with *k*_1_ and *k*_2_ independent input variables, respectively, then the combined BF *u* = *u*_1_ ⊙ *u*_2_ is also unate.

*Proof*: Consider the minimum Boolean expression for the UFs *u*_1_ and *u*_2_. For convenience, let us denote the minimal expressions by the same symbols *u*_1_ and *u*_2_. Since each input variable in *u*_1_ and *u*_2_ is either a positive (*x_i_*) or a negative 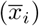 literal (see Property IV.2), the combined expression *u* = *u*_1_ ⊙ *u*_2_ composed of (*k*_1_ + *k*_2_) distinct input variables (due to independence) will also have each variable occur as only its positive or negative literal. This implies that the combined function *u* is a UF. As an example, consider the UFs 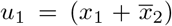 and 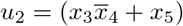. The combined BF *u* = *u*_1_ ⊙ *u*_2_, under the AND operation is simply, 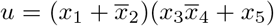. Since each literal appears in *u* only in its positive or its negative form in the expression for *u*, the combined BF is UF.

#### Property B.2.

If a BF *f* is not unate, then the minimal expression for the BF contains both positive and negative literals for at least one of the input variables.

For illustration, consider two BFs 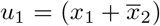 and 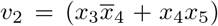, where *u*_1_ is unate while *v*_2_ is not unate. The combined BF 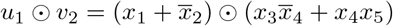 is also not unate, since *x*_4_ occurs as both a positive and a negative literal.

### Nested Canalyzing functions

#### Property B.3.

NCFs have odd bias.

*Proof*: Consider the base case of a NCF with 1 input with the representative expression NCF (1) = *x*_1_. Clearly, the Boolean expression *f* = *x*_1_ refers to the BFs with output vector (0, 1) or (1, 0), both of which have odd bias. Next, let us hypothesize that the NCF with *k* inputs, i.e., *NCF*(*k*), has odd bias *P*. We can then proceed by induction. The NCF with *k* + 1 inputs is given by *NCF*(*k* + 1) = *x*_*k*+1_ ⊙ *NCF*(*k*) by definition (Eq. 8). Since the two independent BFs *x*_*k*+1_ and *NCF*(*k*) have odd bias, using Property A.3, the combined BF *NCF*(*k*+1) will also have an odd bias. Note that a different proof for this property of NCFs was provided by Nikolajewa *et al*. [50].

#### Property B.4.

NCFs are EFs.

*Proof*: Since BFs with odd bias are EFs, using Properties V.1 and B.3, NCFs are also EFs.

#### Property B.5.

NCFs are UFs [30].

*Proof*: Following Aracena [30], since each variable or literal in the expression for NCF (Eq. 8) appears exactly once, it follows that each variable is fixed to either its positive or negative form in the function’s canonical NCF form. Thus, NCFs are UFs using Property IV.2.

### Read-once functions

#### Property B.6.

RoFs are EFs.

*Proof*: From Property V.2, RoFs have odd bias. From Property V.1, BFs with odd bias are EFs. Thus, RoFs are EFs.

#### Property B.7.

RoFs are UFs.

*Proof*: Since each variable or literal in the expression for a RoF (Eq. 9) appears exactly once, it follows that each variable is fixed to either its positive or negative form in the RoF logical expression. Thus, RoFs are UFs according to Property IV.2.

#### Property B.8.

All NCFs are RoFs but not all RoFs are NCFs.

*Proof*: It is evident from Eq. 9 for RoFs and Eq. 8 for NCFs that NCFs are a subset of RoFs. We refer to RoFs which are not NCFs as ‘non-NCF RoFs’.

For number of inputs *k* = 1, 2 and 3, we find that every RoF is a NCF using Properties B.9, B.10 and B.11. For number of inputs *k* ≥ 4, we find that a RoF can be a non-NCF RoF. For example, the function *f*_5_ = *x*_1_*x*_2_ + *x*_3_*x*_4_ is a non-NCF RoF among RoFs with *k* = 4.

#### Property B.9.

RoFs with bias *P* equal to 1 are NCFs.

*Proof*: The DNF of a BF with *k* inputs and bias *P* equal to 1 has just one term, the conjunction of *k* literals, and thus, the function is a NCF. It also follows that RoF with bias *P* equal to 2^*k*^ – 1, which is a complement of the RoF with bias *P* equal to 1, is also a NCF.

#### Property B.10.

RoFs with bias *P* equal to 3 are NCFs.

*Proof*: For *k* = 1, there are no BFs with bias *P* equal to 3. For *k* = 2, RoFs with bias *P* = 1, and their complements with bias *P* = 3, are NCFs using Property B.9. The proof for RoFs with *k* > 2 can be divided into two parts.

Firstly, we show that it is impossible to have a RoF with *k* > 2 and bias *P* equal to 3 by combining RoFs with the OR operator. Let *P*_OR_ be the bias of RoF_OR_(*k* > 2) = *RoF*(*k*_1_) ∨ *RoF*(*k*_2_), such that *k* = *k*_1_ + *k*_2_. Further, let *P*_1_ and *P*_2_ be the biases of *RoF*(*k*_1_) and *RoF*(*k*_2_), respectively. From Property A.2, we have *P_OR_* = 2^*k*_1_^ *P*_2_ + 2^*k*_2_^ *P*_1_ – *P*_1_*P*_2_, which is an increasing monotonic function of *P*_1_ and *P*_2_. This increasing monotonic behaviour can be checked easily by fixing one variable and varying the other. The minimum value of *P*_OR_, *min*(*P*_OR_), is therefore obtained at *P*_1_ = *min*(*P*_1_) = 1, *P*_2_ = *min*(*P*_2_) = 1. Thus, *min*(*P*_OR_) = 2^*k*_1_^ + 2^*k*_2_^ – 1. If *k*_1_ or *k*_2_ is greater than 1, this will result in *min*(*P*_OR_) > 3. As *k* = *k*_1_ + *k*_2_, for *k* > 2, we have *min*(*P*_OR_) > 3. Thus, the bias of *RoF*_OR_ for *k* > 2 cannot be 3.

Secondly, it follows that a RoF with *k* > 2 and bias *P* equal to 3 can only be generated by combining RoFs with the AND operator. Let *P*_AND_ be the bias of *RoF*_AND_(*k*) = *RoF*(*k*_1_) ∧ *RoF*(*k*_2_), such that *k* = *k*_1_ + *k*_2_. From Property A.1, we have *P_AND_* = *P*_1_*P*_2_. Since 3 is prime, a *RoF_AND_*(*k*) with bias 3 can be generated only by combining two RoFs, *RoF*(*k*_1_) and *RoF*(*k*_2_), with biases 1 and 3 (the order being unimportant). For the sake of clarity, let *RoF*(*k*_2_) have bias 3. This would imply that *RoF*(*k*_2_) would in turn have to be generated by combining two RoFs, *RoF*(*k*_2,1_) and *RoF*(*k*_2,2_), with biases 1 and 3, respectively, and so on. Proceeding in this manner, we will be left with a *nested* RoF, with exactly one term having bias 3, and all other RoFs in the *nested* expression having bias 1. In other words, the *nested* RoF would be of the form *x*_1_∧*x*_2_∧*x*_3_∧…∧*x*_*k*−2_∧(*x*_*k*−1_∨*x_k_*), which is a NCF (see Eq. 8). Moreover, it follows that any RoF with bias *P* equal to 2^*k*^ – 3, which is a complement of a RoF with bias 3, is also a NCF.

#### Property B.11.

RoFs with bias *P* equal to 5 are NCFs.

*Proof*: For *k* = 1 and *k* = 2, there are no BFs with bias *P* equal to 5. For *k* = 3, RoFs with bias *P* = 3, and their complements with bias *P* = 5, are NCFs using Property B.10. The proof for RoFs with *k* > 3 can be divided into two parts.

Firstly, we show that it is impossible to have a RoF with *k* > 3 and bias *P* equal to 5 by combining RoFs with the OR operator. Let *P*_OR_ be the bias of *RoF*_OR_(*k* > 3) = *RoF*(*k*_1_) ∨ *RoF*(*k*_2_), such that *k* = *k*_1_ + *k*_2_. From Property A.2, we have *P_OR_* = 2^*k*_1_^ *P*_2_ + 2^*k*_2_^*P*_1_ – *P*_1_*P*_2_, which is an increasing monotonic function of *P*_1_ and *P*_2_. The minimum value of *P*_OR_, *min*(*P*_OR_), is therefore obtained at *P*_1_ = *min*(*P*_1_) = 1, *P*_2_ = *min*(*P*_2_) = 1. Thus, *min*(*P*_OR_) = 2^*k*_1_^ + 2^*k*_2_^ – 1. As *k* = *k*_1_ + *k*_2_, for *k* > 3, we have *min*(*P*_OR_) > 5. Thus, the bias of *RoF*_OR_ for *k* > 3 cannot be 5.

Secondly, it follows from the above that RoF with *k* > 3 and bias *P* equal to 5 can only be generated by combining RoFs with the AND operator. Let *P*_AND_ be the bias of *RoF*_AND_(*k*) = *RoF*(*k*_1_) ∧ *RoF*(*k*_2_), such that *k* = *k*_1_ + *k*_2_. From Property A.1, we have *P*_AND_ = *P*_1_*P*_2_. Since 5 is prime, a *RoF*_AND_(*k*) with bias 5 can be generated only by combining two RoFs, *RoF*(*k*_1_) and *RoF*(*k*_2_), with biases 1 and 5 (the order being unimportant). For the sake of clarity, let *RoF*(*k*_2_) have bias 5. This would imply that *RoF*(*k*_2_) would in turn have to be generated by combining two RoFs, *RoF*(*k*_2,1_) and *RoF*(*k*_2,2_), with biases 1 and 5, respectively, and so on. Proceeding in this manner, we will be left with a *nested* RoF, with exactly one term having bias 5, and all other RoFs in the *nested* expression having bias 1. In other words, the *nested* RoF would be of the form *x*_1_ ∧ *x*_2_ ∧ *x*_3_ ∧ … ∧ *x*_*k*−3_ ∧ (*x*_*k*−2_ ∨ (*x*_*k*−1_ ∧ *x_k_*)), which is a NCF (see Eq. 8). Moreover, it follows that any RoF with bias *P* equal to 2^*k*^ – 5, which is a complement of an RoF with bias 5, is also a NCF.

In Fig. 4, we find that when a NCF and a non-NCF RoF belong to the same *k*[*P*] set, then the NCF has a lower average sensitivity compared to the non-NCF RoF. Further, we have shown that NCFs have the *minimum* average sensitivity in any *k*[*P*] set with odd bias *P*. We were then curious as to whether 2 representative non-NCF RoFs within a *k*[*P*] set could have the same average sensitivity, and we find this is indeed true via exhaustive computational enumeration of RoFs with *k* ≤ 10. We observe that such a case of 2 representative non-NCF RoFs first occur when *k* = 7 at the bias *P* = 25. In Appendix E, we describe our program to check if a user-specified BF is a RoF.

## Appendix C: Biological dataset compiling Boolean functions from reconstructed discrete models of living systems

In order to assess the abundance of different types of biologically meaningful BFs in reconstructed discrete models of living systems, we first compiled a large dataset of 88 models which have been published to date. These 88 models were either downloaded from databases such as Cell Collective [25] (https://cellcollective.org/), GIN-SIM [58] (http://ginsim.org/) or BioModels [59] (http://www.ebi.ac.uk/biomodels/), or directly obtained from the corresponding published article. Notably, most of these 88 models were downloaded from the Cell collective database [25]. Further, this compilation of 88 models spans the overwhelming majority of Boolean models of biological systems reconstructed and published to date. The majority of these models pertains to mammalian systems and a much smaller fraction pertains to plant systems. The mammalian models include networks for signaling pathways [24, 64, 65], differentiation [66, 67] and various cancers [68–70]. Among the plant models, this compilation includes cases from flower organ specification [67], root stem cells [71], and guard cell signalling [72]. Overall, the 88 discrete models in this compilation capture a very diverse collection of biological processes throughout multiple kingdoms of life.

This study is focused only on properties of BFs assigned to different nodes in reconstructed models of biological networks. Some of those networks included nodes taking more than two discrete states; in our compilation, we included only BFs assigned to nodes with binary states which further also had inputs only from other nodes with binary states. While compiling the BFs from these 88 models, we have also gathered the information on the *signs* of interactions between regulators (input nodes) and target gene (output node). Such information is typically obtained from associated experimental literature.

Across the 88 models in this compilation, the number of nodes in a model varies between 4 and 128. From these 88 models, we have compiled 2687 BFs pertaining to 2687 nodes that have number of inputs *k* ≥ 1 (Fig. 3(b)). The BFs assigned to each node in these 88 models are the result of many authors manually identifying appropriate input-output relations during network reconstruction. In other words, the 2687 BFs in the reference biological dataset were chosen during model reconstruction process such as to capture the known regulatory information.

## Appendix D: Statistical Tests

### Enrichments and relative enrichments

Consider a given type of BF (say unate with *k* inputs) which we refer to as *T*. Denote by *f*_0_ the fraction of functions that are of type T in the random ensemble and by *f*_1_ the corresponding fraction in our reference biological dataset. The enrichment ratio E is simply *f*_1_/*f*_0_. If *E* > 1, then the *T* is enriched while if *E* < 1, *T* is depleted. If *E* = 1, there is neither enrichment nor depletion.

In our study we are also interested in relative enrichments to probe for possible causes of enrichments. For that we consider a type *T* and one of its subtypes, say *T_s_*. For instance in our comparison of the two measures of complexity we examined the case where *T*=RoF and *T_s_*=NCF. In direct analogy with what was done for enrichments, we define the relative enrichment *E_R_* = (*f*_*s*,1_/*f*_1_)/(*f*_*s*,0_/*f*_0_) where the subscript *s* refers to type *T_s_*. If enrichment is driven solely by the property of being in *T*, then the relative enrichment is expected to be close to 1. As a consequence, if *E_R_* is large, then there must be other factors than “belonging to *T*” driving this relative enrichment.

### Associated p-values

We developed a first statistical test to determine whether an observed enrichment *E* was statistically significant. The underlying statistical distribution of the random variable *E* is obtained by formalizing an underlying hypothesis referred to as *H*_0_. Here *H*_0_ corresponds to hypothesizing that the functions in the reference biological dataset are drawn from the random ensemble where all 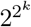 BFs with *k* inputs are equiprobable. The (right-sided) p-value is then just the probability that such a drawing leads to a value of E as large as the one actually observed. This probability is computed as follows. The fraction *f*_0_ is first determined (see Table S2). Then we consider drawing *M* BFs from the random ensemble and count the number *m* of these functions that belong to type *T* (*M* is the number of BFs in the reference biological dataset). The probability of having a given value *m* is given by the binomial distribution: 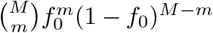. The desired p-value is then just the sum of all such probabilities under the condition that *m* is larger or equal to *M f*_1_. This sum is computed numerically.

The second type of test we perform concerns the statistical significance of a relative enrichment *E_R_* deviating from 1. Again we formalize this by introducing an *H*_0_ hypothesis. Using the notation of the previous subsection, *H*_0_ corresponds to assuming that although there is a selection for *T* (as evident from a large value of *E*), the elements that are drawn within *T* have a uniform probability, that is members of *T_s_* are not more probable than the other elements of *T*. Consider then drawing a sample of size *M* under *H*_0_. If it leads to *M_T_* elements in *T* as in the reference biological dataset, the distribution of the number of elements in *T_s_* is known. Specifically, the probability to have m elements in *T_s_* is 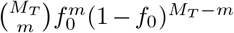 where now *f*_0_ is the ratio of sizes of *T_s_* and *T* in the random ensemble of *T_s_* (see Table S2). The desired p-value is then just the sum of all such probabilities under the condition that *m* is larger or equal to the number of *T_s_* elements in the reference biological dataset. Again, this sum is computed numerically.

The number of functions belonging to a particular type of BF was obtained from both computation and theory. The number of CFs for *k* = 6, 7, 8 and NCFs for *k* = 7, 8 were obtained from [73] and [74]. In certain cases (EFs and UFs having 6, 7 or 8 inputs), it was computationally unfeasible to obtain the exact number of functions in these types and there was no data in the literature as well, hence we used sampling to estimate the probability of a BF to belong to these types, for the specified number of inputs.

## Appendix E: RoF checker

To check whether a BF is a RoF, we make use of the various properties of RoFs. To begin with, we generate a representative RoF for each equivalence class, going up to *k* =10 inputs using the Property IV.8. We store the truth table, bias and the average sensitivity of each representative RoF in computer memory so that it can be used as a lookup table. Next, we implement the procedure shown in the flowchart (see Fig. S7). This program takes as input a BF via its truth table representation; the bias of the BF is determined. The program then proceeds by performing successive tests, from quite simple to more complex, as follows. If the bias is even then the BF is not a RoF (see Property V.2). Since NCFs are a subset of RoFs, we check whether the BF is a NCF as that is relatively simple computationally (just successively determine the canalyzing input variables). If the function is not a NCF, we check whether the input BF is a UF since RoFs are unate. If the BF is unate, we calculate average sensitivity. Then we use the lookup table to extract all of the representative RoFs having that bias and average sensitivity. Recall that all elements in an equivalence class have the same bias and average sensitivity. In case no representative RoF matches, then the BF is not a RoF. Assuming that there is at least one representative RoF extracted, the program then loops over that list. For each such RoF we generate all RoFs belonging to that same equivalence class (just loop over all isomorphisms), and for each such function the program directly checks whether its truth table is the same as that of the input BF. If an equality is found, the input BF is a RoF and we are done. If all the representative RoFs are tested without any success, then the input BF is not a RoF. The catalog of RoFs along with the python code to check for RoFs is available via the GiHub repository: https://github.com/asamallab/MCBF.

## ACKNOWLEDGMENTS

A. Subbaroyan would like to thank Sathish Kumar for discussions, and Sneha Subbaroyan for help with figures. A. Samal acknowledges support from the Max Planck Society, Germany, through the award of a Max Planck Partner Group in Mathematical Biology. IPS2 benefits from the support of Saclay Plant Sciences-SPS (ANR-17-EUR-0007).

## SUPPLEMENTAL MATERIAL

**TABLE S1.** The number of Boolean functions (BFs) belonging to the different types, at a given number of inputs *k* ≤ 5. Here, EF corresponds to effective functions, UF to unate functions (all sign combinations), CF to canalyzing functions, EUF to effective and unate functions, ECF to effective and canalyzing functions, UCF to unate and canalyzing functions, EUCF to effective, unate and canalyzing functions. In addition, the table lists the total number of BFs for each *k*.

| $k$ | Types of BFs | | | | | | | |
| --- | --- | --- | --- | --- | --- | --- | --- | --- |
|  | All | EF | UF | CF | EUF | ECF | UCF | EUCF |
| 1 | 4 | 2 | 4 | 4 | 2 | 2 | 4 | 2 |
| 2 | 16 | 10 | 14 | 14 | 8 | 8 | 14 | 8 |
| 3 | 256 | 218 | 104 | 120 | 72 | 88 | 96 | 64 |
| 4 | 65536 | 64594 | 2170 | 3514 | 1824 | 3104 | 1178 | 864 |
| 5 | 4294967296 | 4294642034 | 230540 | 1292276 | 220608 | 1275784 | 36796 | 31744 |

**TABLE S2.** The fraction of Boolean functions (BFs) belonging to the different types, at a given number of inputs *k* ≤ 5. Here, EF corresponds to effective functions, UF to unate functions (all sign combinations), CF to canalyzing functions, EUF to effective and unate functions, ECF to effective and canalyzing functions, UCF to unate and canalyzing functions, EUCF to effective, unate and canalyzing functions.

| $k$ | Types of BFs | | | | | | |
| --- | --- | --- | --- | --- | --- | --- | --- |
|  | EF | UF | CF | EUF | ECF | UCF | EUCF |
| 1 | 0.500 | 1.000 | 1.000 | 0.500 | 0.500 | 1.000 | 0.500 |
| 2 | 0.625 | 0.875 | 0.875 | 0.500 | 0.500 | 0.875 | 0.500 |
| 3 | 0.852 | 0.406 | 0.469 | 0.281 | 0.344 | 0.375 | 0.250 |
| 4 | 0.986 | 0.033 | 0.054 | 0.028 | 0.047 | 0.018 | 0.013 |
| 5 | 1.000 | $5.37 \times 10^{-5}$ | $3.01 \times 10^{-4}$ | $5.14 \times 10^{-5}$ | $2.97 \times 10^{-4}$ | $8.57 \times 10^{-6}$ | $7.39 \times 10^{-6}$ |

**TABLE S3.** Parity distribution of biologically meaningful types of Boolean functions (BFs) for a given number of inputs *k*. The fraction of even parity functions of a particular type of BF is calculated with respect to the total number of functions of that BF type. Here, EF corresponds to effective functions, UF to unate functions (all sign combinations), CF to canalyzing functions, EUF to effective and unate functions, ECF to effective and canalyzing functions, UCF to unate and canalyzing functions, EUCF to effective, unate and canalyzing functions. Note that entries labeled ‘-’ are those which could not be computed due to inadequate computational resources.

| $k$ | Number of even parity functions | | | | | | | | Fraction of even parity functions | | | | | | | |
| --- | --- | --- | --- | --- | --- | --- | --- | --- | --- | --- | --- | --- | --- | --- | --- | --- |
|  | All | EF | UF | CF | EUF | ECF | UCF | EUCF | All | EF | UF | CF | EUF | ECF | UCF | EUCF |
| 1 | 2 | 0 | 2 | 2 | 0 | 0 | 2 | 0 | 0.5 | 0 | 0.5 | 0.5 | 0 | 0 | 0.5 | 0 |
| 2 | 8 | 2 | 6 | 6 | 0 | 0 | 6 | 0 | 0.5 | 0.2 | 0.429 | 0.429 | 0 | 0 | 0.429 | 0 |
| 3 | 128 | 90 | 40 | 56 | 8 | 24 | 32 | 0 | 0.5 | 0.413 | 0.385 | 0.467 | 0.111 | 0.273 | 0.333 | 0 |
| 4 | 32768 | 31826 | 922 | 1754 | 576 | 1344 | 442 | 128 | 0.5 | 0.493 | 0.425 | 0.499 | 0.316 | 0.433 | 0.375 | 0.148 |
| 5 | 2147483648 | - | 110348 | 646132 | 100416 | 629640 | 15932 | 10880 | 0.5 | - | 0.479 | 0.5 | 0.455 | 0.494 | 0.433 | 0.343 |

**TABLE S4.** The number of effective and unate functions (EUFs) at a given number of inputs *k* ≤ 5 for different combinations of activators and inhibitors. The table also gives the fraction of EUFs that have even bias for different sign combinations. These numbers have been obtained via exhaustive computational enumeration of EUFs.

| $k$ | Activators | Inhibitors | EUFs | | |
| --- | --- | --- | --- | --- | --- |
|  |  |  | Total | Even Bias | Fraction with Even bias |
| 1 | 1 | 0 | 1 | 0 | 0 |
|  | 0 | 1 | 1 | 0 | 0 |
| 2 | 2 | 0 | 2 | 0 | 0 |
|  | 1 | 1 | 4 | 0 | 0 |
|  | 0 | 2 | 2 | 0 | 0 |
| 3 | 3 | 0 | 9 | 1 | 0.111 |
|  | 2 | 1 | 27 | 3 | 0.111 |
|  | 1 | 2 | 27 | 3 | 0.111 |
|  | 0 | 3 | 9 | 1 | 0.111 |
| 4 | 4 | 0 | 114 | 36 | 0.316 |
|  | 3 | 1 | 456 | 144 | 0.316 |
|  | 2 | 2 | 684 | 216 | 0.316 |
|  | 1 | 3 | 456 | 144 | 0.316 |
|  | 0 | 4 | 114 | 36 | 0.316 |
| 5 | 5 | 0 | 6894 | 3138 | 0.455 |
|  | 4 | 1 | 34470 | 15690 | 0.455 |
|  | 3 | 2 | 68940 | 31380 | 0.455 |
|  | 2 | 3 | 68940 | 31380 | 0.455 |
|  | 1 | 4 | 34470 | 15690 | 0.455 |
|  | 0 | 5 | 6894 | 3138 | 0.455 |

**TABLE S5.**
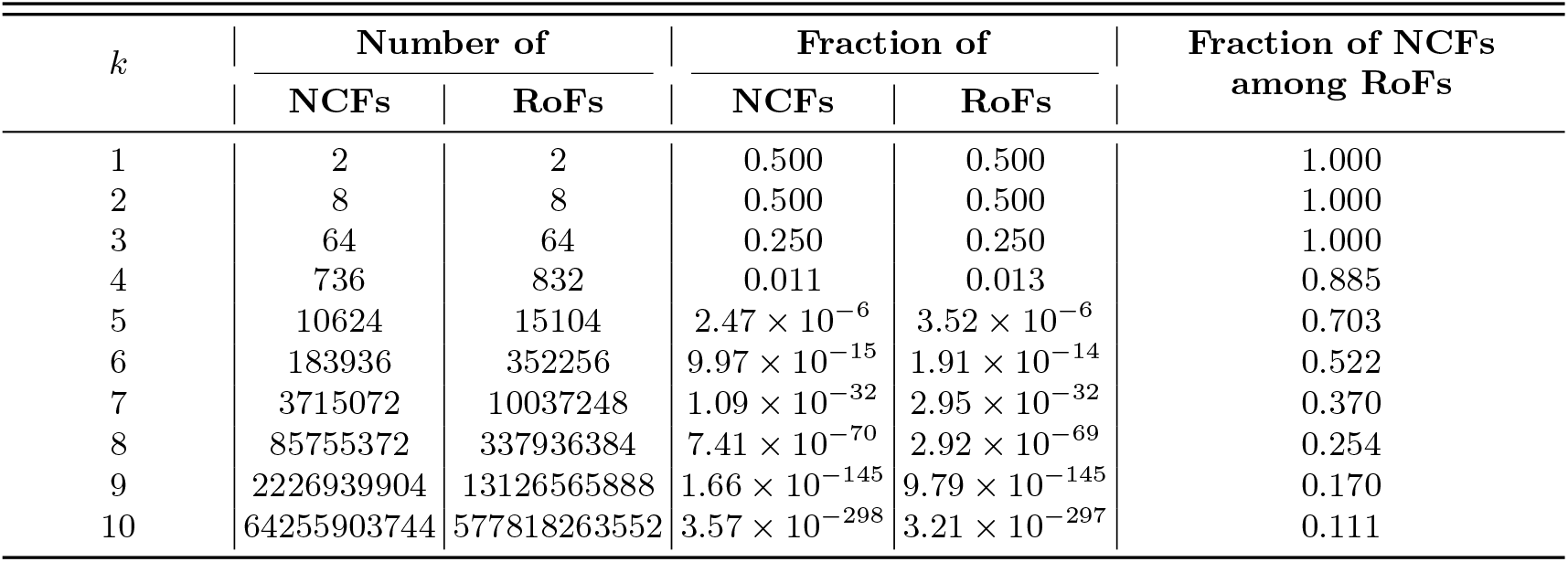
The number of Nested Canalyzing functions (NCFs) and Read-once functions (RoFs), and their fraction among all Boolean functions (BFs) for a given number of inputs *k*. In addition, the table lists the fraction of NCFs among RoFs for a given number of inputs *k*.

**TABLE S6.**
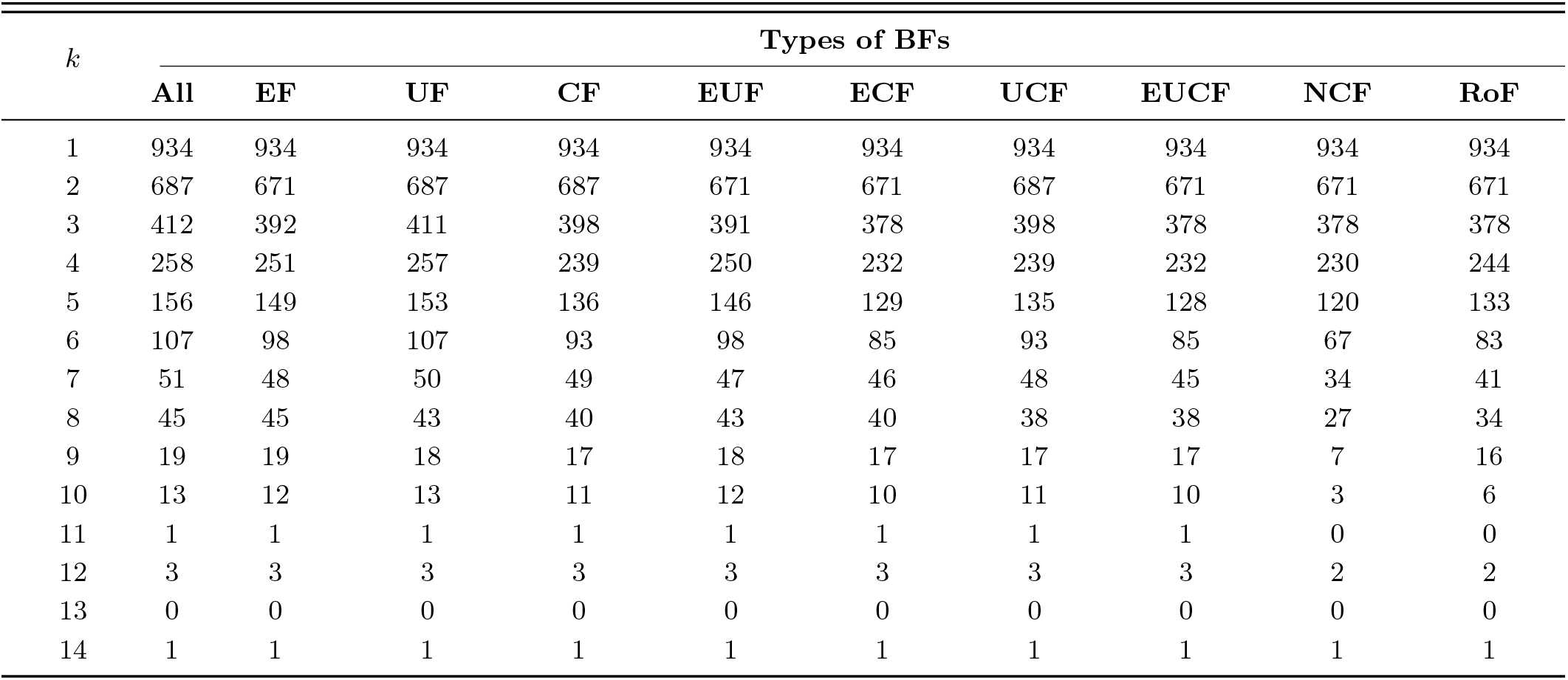
Number of different types of biologically meaningful Boolean functions (BFs) in the reference biological dataset. Here, *k* is the number of inputs, ‘All’ is the total number of BFs for a given number of inputs, EF corresponds to effective functions, UF to unate functions (all sign combinations), CF to canalyzing functions, EUF to effective and unate functions, ECF to effective and canalyzing functions, UCF to unate and canalyzing functions, EUCF to effective, unate and canalyzing functions, NCF to Nested Canalyzing functions and RoF to Read-once functions.

**TABLE S7.** Fraction of different types of biologically meaningful Boolean functions (BFs) in the reference biological dataset. Here, *k* is the number of inputs, EF corresponds to effective functions, UF to unate functions (all sign combinations), CF to canalyzing functions, EUF to effective and unate functions, ECF to effective and canalyzing functions, UCF to unate and canalyzing functions, EUCF to effective, unate and canalyzing functions, NCF to Nested Canalyzing functions and RoF to Read-once functions.

| $k$ | Types of BFs | | | | | | | | |
| --- | --- | --- | --- | --- | --- | --- | --- | --- | --- |
|  | EF | UF | CF | EUF | ECF | UCF | EUCF | NCF | RoF |
| 1 | 1 | 1 | 1 | 1 | 1 | 1 | 1 | 1 | 1 |
| 2 | 0.977 | 1 | 1 | 0.977 | 0.977 | 1 | 0.977 | 0.977 | 0.977 |
| 3 | 0.951 | 0.998 | 0.966 | 0.949 | 0.917 | 0.966 | 0.917 | 0.917 | 0.917 |
| 4 | 0.973 | 0.996 | 0.926 | 0.969 | 0.899 | 0.926 | 0.899 | 0.891 | 0.946 |
| 5 | 0.955 | 0.981 | 0.872 | 0.936 | 0.827 | 0.865 | 0.821 | 0.769 | 0.853 |
| 6 | 0.916 | 1 | 0.869 | 0.916 | 0.794 | 0.869 | 0.794 | 0.626 | 0.776 |
| 7 | 0.941 | 0.980 | 0.961 | 0.922 | 0.902 | 0.941 | 0.882 | 0.667 | 0.804 |
| 8 | 1 | 0.956 | 0.889 | 0.956 | 0.889 | 0.844 | 0.844 | 0.6 | 0.756 |
| 9 | 1 | 0.947 | 0.895 | 0.947 | 0.895 | 0.895 | 0.895 | 0.368 | 0.842 |
| 10 | 0.923 | 1 | 0.846 | 0.923 | 0.769 | 0.846 | 0.769 | 0.231 | 0.462 |
| 11 | 1 | 1 | 1 | 1 | 1 | 1 | 1 | 0 | 0 |
| 12 | 1 | 1 | 1 | 1 | 1 | 1 | 1 | 0.667 | 0.667 |
| 13 | - | - | - | - | - | - | - | - | - |
| 14 | 1 | 1 | 1 | 1 | 1 | 1 | 1 | 1 | 1 |

**TABLE S8.** *p*-value tests for statistical enrichments of the different types of Boolean functions (BFs) in the reference biological dataset. A low p-value indicates that the corresponding type of BF is enriched in the reference biological dataset when compared to the ensemble of all BFs. For *k* > 2 when the p-value shown is 0, it was smaller than what we could measure, and when the p-value shown is 1, its deviation from 1 was smaller than we could measure. Here, EF corresponds to effective functions, UF to unate functions (all sign combinations), CF to canalyzing functions, NCF to Nested Canalyzing functions and RoF to Read-once functions.

| $k$ | Odd bias | EF | UF | CF | NCF | RoF |
| --- | --- | --- | --- | --- | --- | --- |
| 2 | $3.742 \times 10^{-177}$ | $6.75 \times 10^{-114}$ | 0 | 0 | $3.74 \times 10^{-177}$ | $3.74 \times 10^{-177}$ |
| 3 | $6.253 \times 10^{-76}$ | $2.44 \times 10^{-11}$ | $6.65 \times 10^{-162}$ | $1.83 \times 10^{-111}$ | $3.09 \times 10^{-184}$ | $3.09 \times 10^{-184}$ |
| 4 | $2.714 \times 10^{-62}$ | 0.919 | 0 | $8.86 \times 10^{-279}$ | 0 | 0 |
| 5 | $2.718 \times 10^{-25}$ | 1 | 0 | 0 | 0 | 0 |
| 6 | $1.753 \times 10^{-13}$ | 1 | 0 | 0 | 0 | 0 |
| 7 | $6.058 \times 10^{-08}$ | 1 | 0 | 0 | 0 | 0 |
| 8 | $1.561 \times 10^{-06}$ | 1 | 0 | 0 | 0 | 0 |

**TABLE S9.** The enrichment ratios *E_R_* for the RoFs and NCFs in the ensemble of odd bias BFs, EFs and UFs. These ratios indicate the extent of the over-representation of such functions in the reference biological dataset. *E_R_* > 1 suggests that there is indeed an enrichment of RoFs and NCFs within the EFs, UFs and CFs in the reference biological dataset when compared to that expected in the ensemble of all EFs, UFs and CFs.

| $k$ | $E_R$ for RoF in: | | | $E_R$ for NCF in: | | |
| --- | --- | --- | --- | --- | --- | --- |
|  | Odd bias | EF | UF | Odd bias | EF | UF |
| 1 | 1 | 1 | 2 | 1 | 1 | 2 |
| 2 | 1 | 1.25 | 1.709 | 1 | 1.250 | 1.709 |
| 3 | 2 | 3.285 | 1.495 | 2 | 3.285 | 1.495 |
| 4 | 41.29 | 80.421 | 2.476 | 38.749 | 75.472 | 2.334 |
| 5 | $1.76 \times 10^6$ | $3.26 \times 10^5$ | 13.268 | $1.37 \times 10^5$ | $2.54 \times 10^5$ | 11.971 |

**TABLE S10.**
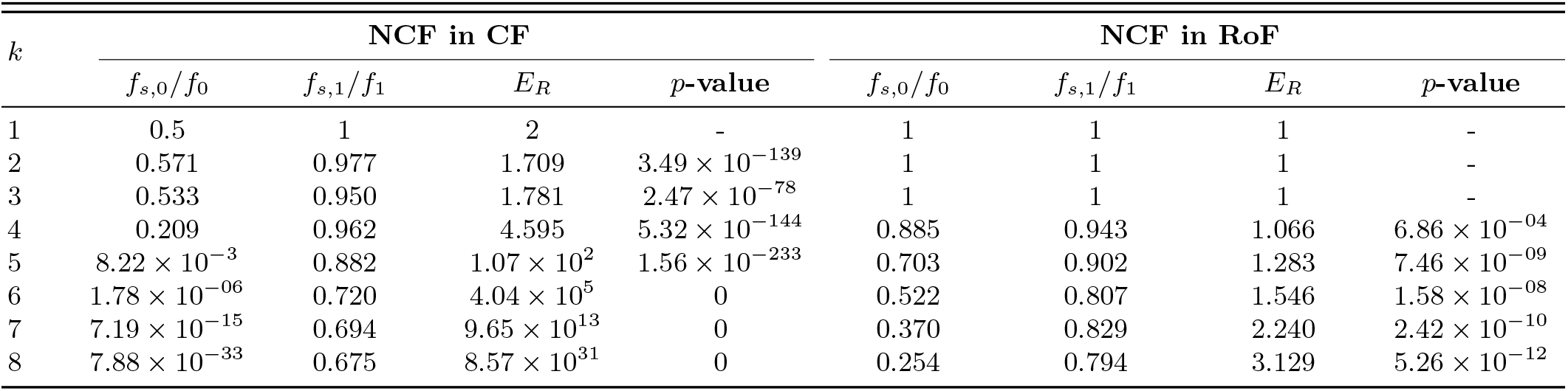
The relative enrichment ratio *E_R_* of the NCFs in the CFs and RoFs. *f*_*s*,0_/*f*_0_ denotes the fractions of functions that are NCFs in the space of all CFs or RoFs and *f*_*s*,1_/*f*_1_, the equivalent fraction in the reference biological dataset. *E_R_* = (*f*_*s*,1_/*f*_1_)/(*f*_*s*,0_/*f*_0_) denotes the enrichment ratio and it indicates the extent of the over-representation of such functions in the reference dataset. Computations are reported for BFs with *k* ≤ 8 inputs. The low p-values indicate that there is an enrichment of NCFs within the CFs and RoFs in the reference dataset when compared to that expected in the ensemble of all CFs and RoFs.

| $k$ | NCF in CF | | | | NCF in RoF | | | |
| --- | --- | --- | --- | --- | --- | --- | --- | --- |
| | $f_{s,0}/f_0$ | $f_{s,1}/f_1$ | $E_R$ | $p$-value | $f_{s,0}/f_0$ | $f_{s,1}/f_1$ | $E_R$ | $p$-value |
| 1 | 0.5 | 1 | 2 | - | 1 | 1 | 1 | - |
| 2 | 0.571 | 0.977 | 1.709 | $3.49 \times 10^{-139}$ | 1 | 1 | 1 | - |
| 3 | 0.533 | 0.950 | 1.781 | $2.47 \times 10^{-78}$ | 1 | 1 | 1 | - |
| 4 | 0.209 | 0.962 | 4.595 | $5.32 \times 10^{-144}$ | 0.885 | 0.943 | 1.066 | $6.86 \times 10^{-04}$ |
| 5 | $8.22 \times 10^{-3}$ | 0.882 | $1.07 \times 10^2$ | $1.56 \times 10^{-233}$ | 0.703 | 0.902 | 1.283 | $7.46 \times 10^{-09}$ |
| 6 | $1.78 \times 10^{-06}$ | 0.720 | $4.04 \times 10^5$ | 0 | 0.522 | 0.807 | 1.546 | $1.58 \times 10^{-08}$ |
| 7 | $7.19 \times 10^{-15}$ | 0.694 | $9.65 \times 10^{13}$ | 0 | 0.370 | 0.829 | 2.240 | $2.42 \times 10^{-10}$ |
| 8 | $7.88 \times 10^{-33}$ | 0.675 | $8.57 \times 10^{31}$ | 0 | 0.254 | 0.794 | 3.129 | $5.26 \times 10^{-12}$ |

**FIG. S1.**
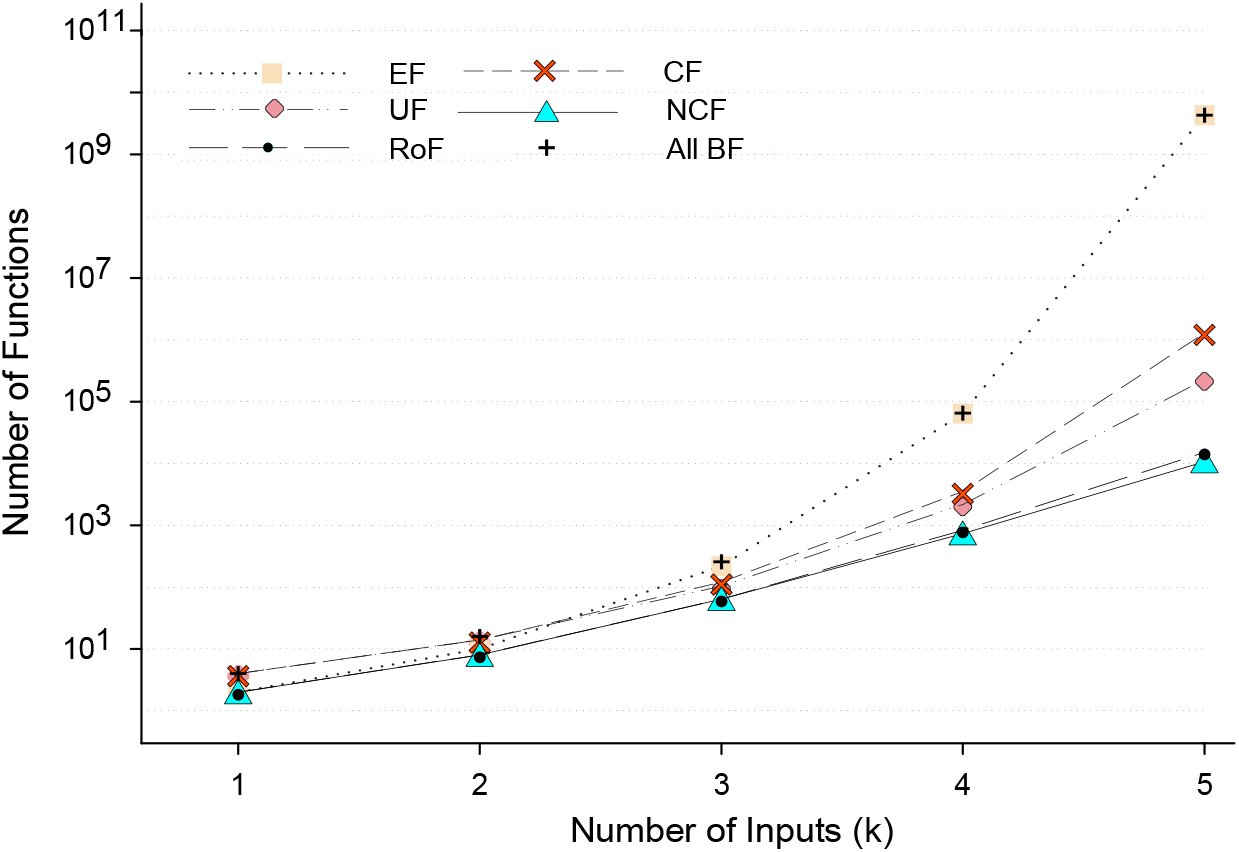
The number of Boolean functions (BFs) for each biologically meaningful type, at a given number of inputs *k* ≤ 5. Here, EF corresponds to effective functions, UF to unate functions (all sign combinations), CF to canalyzing functions, NCF to nested canalyzing functions, RoF to read-once functions, and “All BF” to all 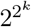 possible functions with *k* inputs.

**FIG. S2.**
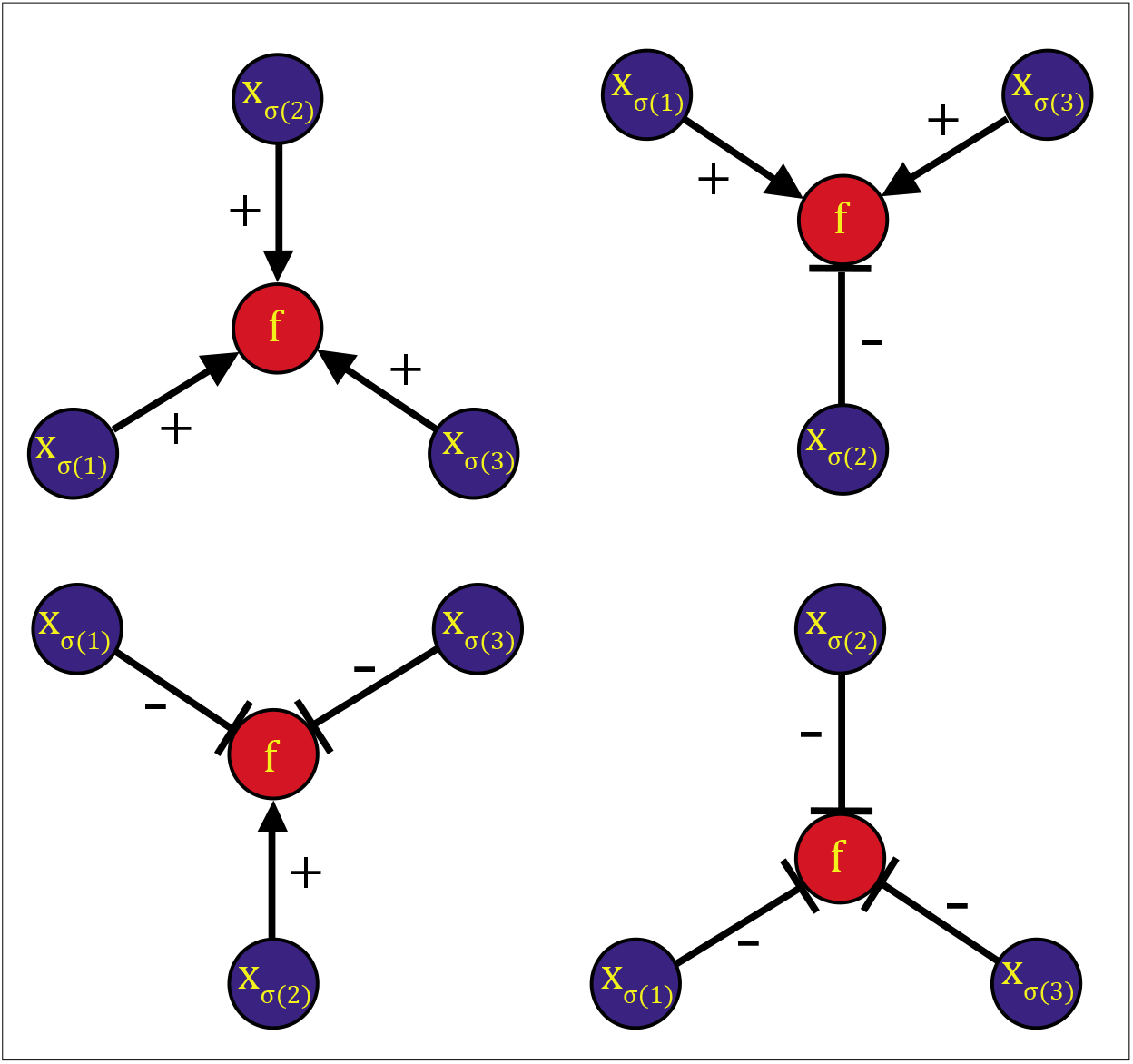
Schematic figure depicting the four possible *sign* combinations for the *k* = 3 inputs to a node (gene) *f*. Each input or regulatory interaction to the node *f* can be an activator or inhibitor which are shown as ‘+’ or ‘−’, respectively. The 3 inputs to the node *f* are labeled by variables *x*_*σ*(1)_, *x*_*σ*(2)_ or *x*_*σ*(3)_, without repetition of any label. *σ* is the permutation of the set {1, 2, 3}, and *σ*(*i*) represents the *i*^th^ element of a given permutation.

**FIG. S3.**
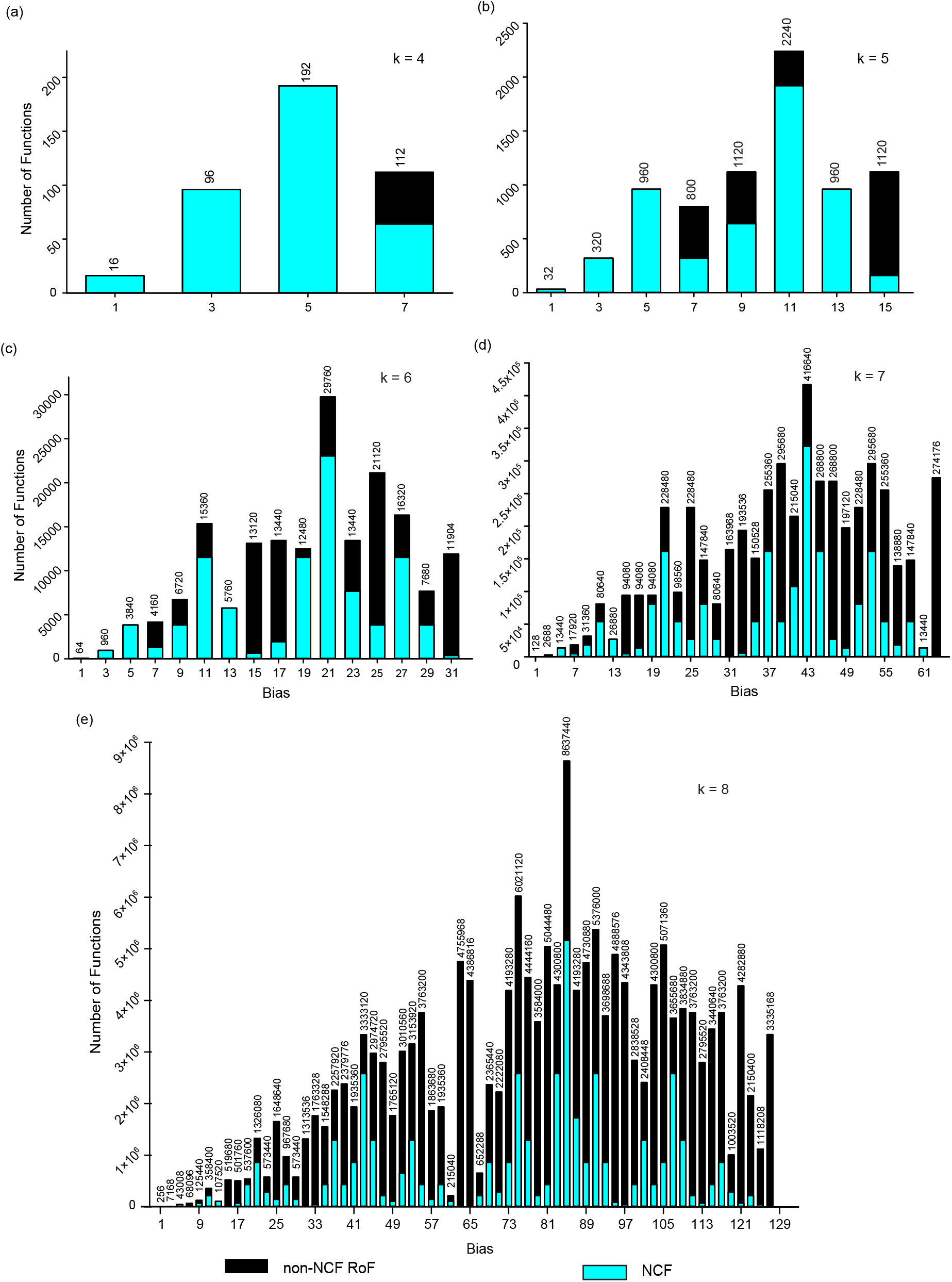
Frequency distribution of read-once functions (RoFs) across different bias *P* for functions with **(a)** *k* = 4, **(b)** *k* = 5, **(c)** *k* = 6, **(d)** *k* = 7, and **(e)** *k* = 8 inputs. For each bar, we display the number of RoFs with that bias value. The figure also gives the frequency distribution of nested canalyzing functions (NCFs), which are a subset of RoFs. Due to the *complementarity* property of Boolean functions, the distribution is symmetric about the bias value 2^*k*−1^ for a given *k*, and therefore, we display only the first half of the distribution, from 0 to 2^*k*−1^ in each case.

**FIG. S4.**
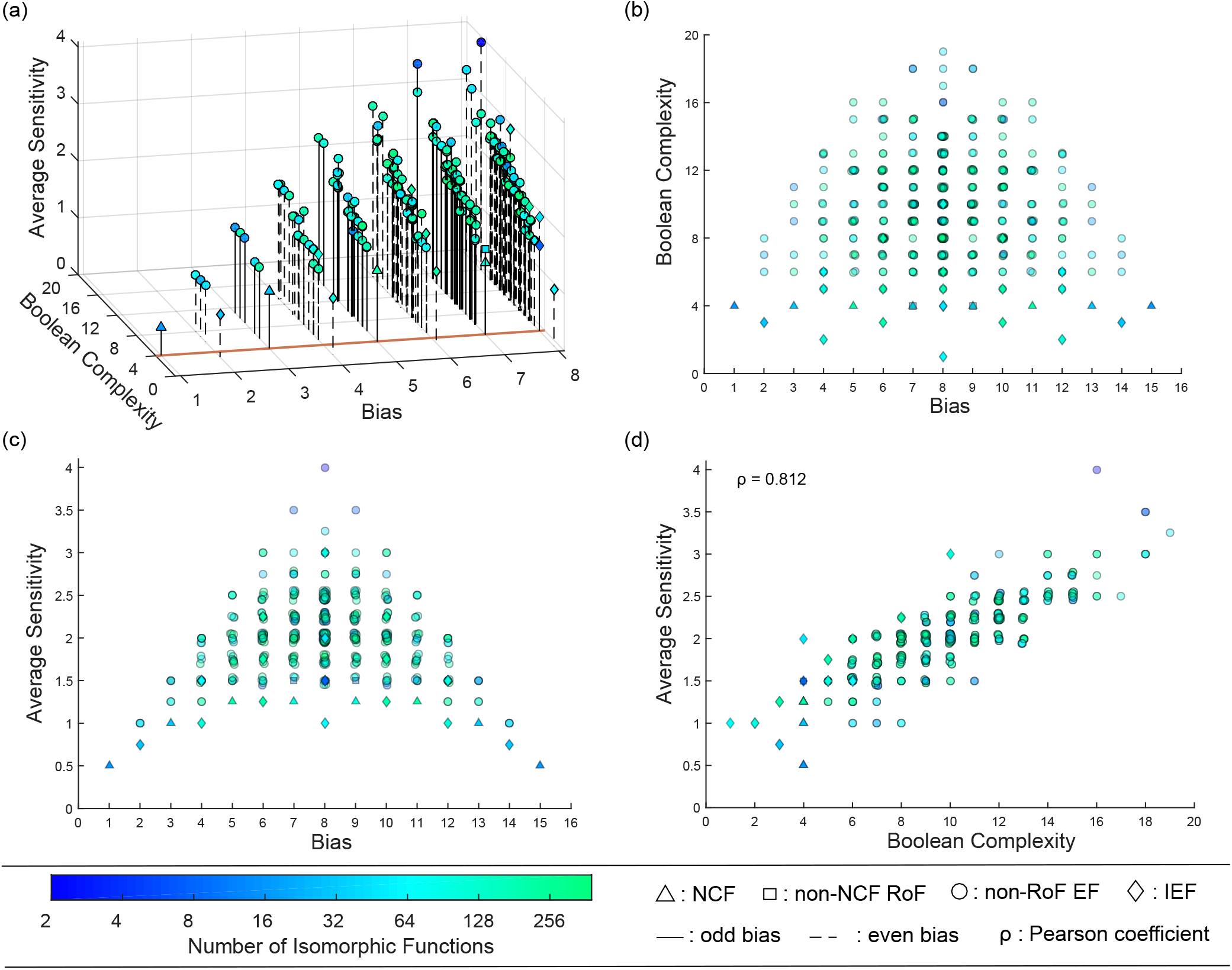
Dependence of the two complexity measures on the bias and associated 2D projections for all Boolean functions (BFs) with *k* = 4 inputs. In each sub-figure, a point (with symbols circle, triangle, square or diamond) is displayed for each representative BF of its class (see Section IIC). The slight ‘jiggle’ at some points is added to resolve the overlapping representative BFs. In this plot, the class ‘non-RoF EF’ refers to the subset of EFs which are not RoFs. The same color scheme for the number of isomorphic functions is applicable to all the plots. **(a)** In this plot, with increasing bias, it is clear that both measures of complexity increase up to *P* = 8. The brown line drawn at the Boolean complexity 4 highlights the functions that possess the minimum Boolean complexity and are effective as well. The NCFs and RoFs are the only functions which lie along this line. Since these complexity measures are invariant under complementation of the BF, the bias values have been shown only up to *P* = 8. **(b)** Variation of the Boolean complexity with increasing bias. With increasing bias, the number of representative BFs increases, but so does the range of Boolean complexity of these functions. The RoFs and ineffective functions (IEFs) have the minimum Boolean complexity in any 4[*P*] set. **(c)** Variation of average sensitivity with increasing bias. Clearly, the NCFs and IEFs have the minimum average sensitivity in any 4[*P*] set. Note that both sub-figures (b) and (c) are symmetric about *P* = 8 due to the complementarity property. **(d)** Correlation of the Boolean complexity and the average sensitivity. The Pearson correlation coefficient (*ρ*) between the two measures was calculated for all BFs with *k* = 4 and *P* ≤ 8. The high Pearson coefficient suggests that there is a positive monotonic relationship between these 2 measures.

**FIG. S5.**
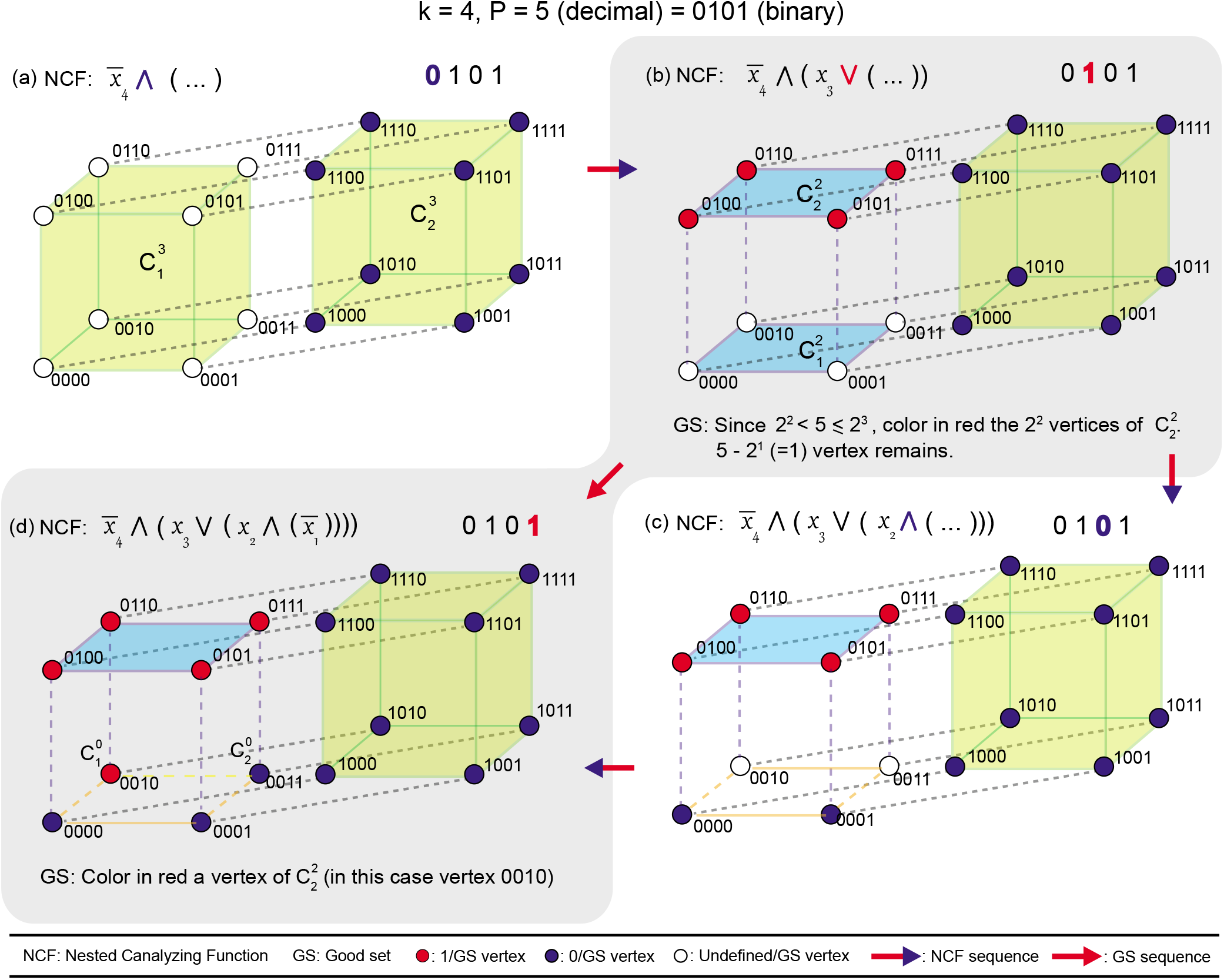
‘Good set’ (GS) for *P* vertices where *P* is odd on a *k*-dimensional hypercube is equivalent to a Nested Canalyzing Function (NCF) in that *k*[*P*] set. In parts (b) and (d) shaded in grey, we show the recursive construction of a GS for *P* = 5 vertices in a 4-dimensional hypercube by coloring it’s vertices red, and in parts (a), (b), (c) and (d), we show the equivalence of that GS of 5 vertices to a NCF with bias 5. The vertices of the hypercube are labeled in the order *x*_4_, *x*_3_, *x*_2_, *x*_1_ wherein *x_i_* is 0 or 1. Here, 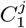 and 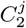 denote the two vertex disjoint *j*-dimensional hypercubes of the (*j* + 1)-dimensional hypercube. The ‘active’ bit in each part (a), (b), (c) and (d) is the colored bit in the binary representation of 5 in that part. **(a)** The vertices with *x*_4_ = 1 are canalyzed to the output value 0. The active bit in this step is 0 and as a result the ∧ operator follows the literal 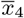. **(b)** Since *P* = 5 lies between 2^2^ and 2^3^, 2^2^ vertices of either 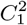 or 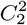 (here, 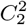) form part of the GS. This leaves 5 – 4 = 1 vertex to be colored red to complete the GS. This choice of 4 vertices in 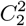 for the GS leads to the canalyzation of vertices labelled *x*_4_ = 0 and *x*_3_ = 1 to the output value 1. The active bit in this step is 1 and as a result the ∨ operator follows the literal *x*_3_. **(c)** The vertices with *x*_4_ = 0, *x*_3_ = 0 and *x*_2_ =0 are canalyzed to the output value 0. The active bit in this step is 0 and as a result the ∨ operator follows the literal *x*_2_. **(d)** For the last step, any vertex in 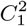 can be colored to complete the 5 vertices in GS, and we color the vertex 0010. The vertex with *x*_4_ = 0, *x*_3_ = 0, *x*_2_ = 1 and *x*_4_ = 0 is canalyzed to the output value 1, and the one remaining vertex is set to output value 0.

**FIG. S6.**
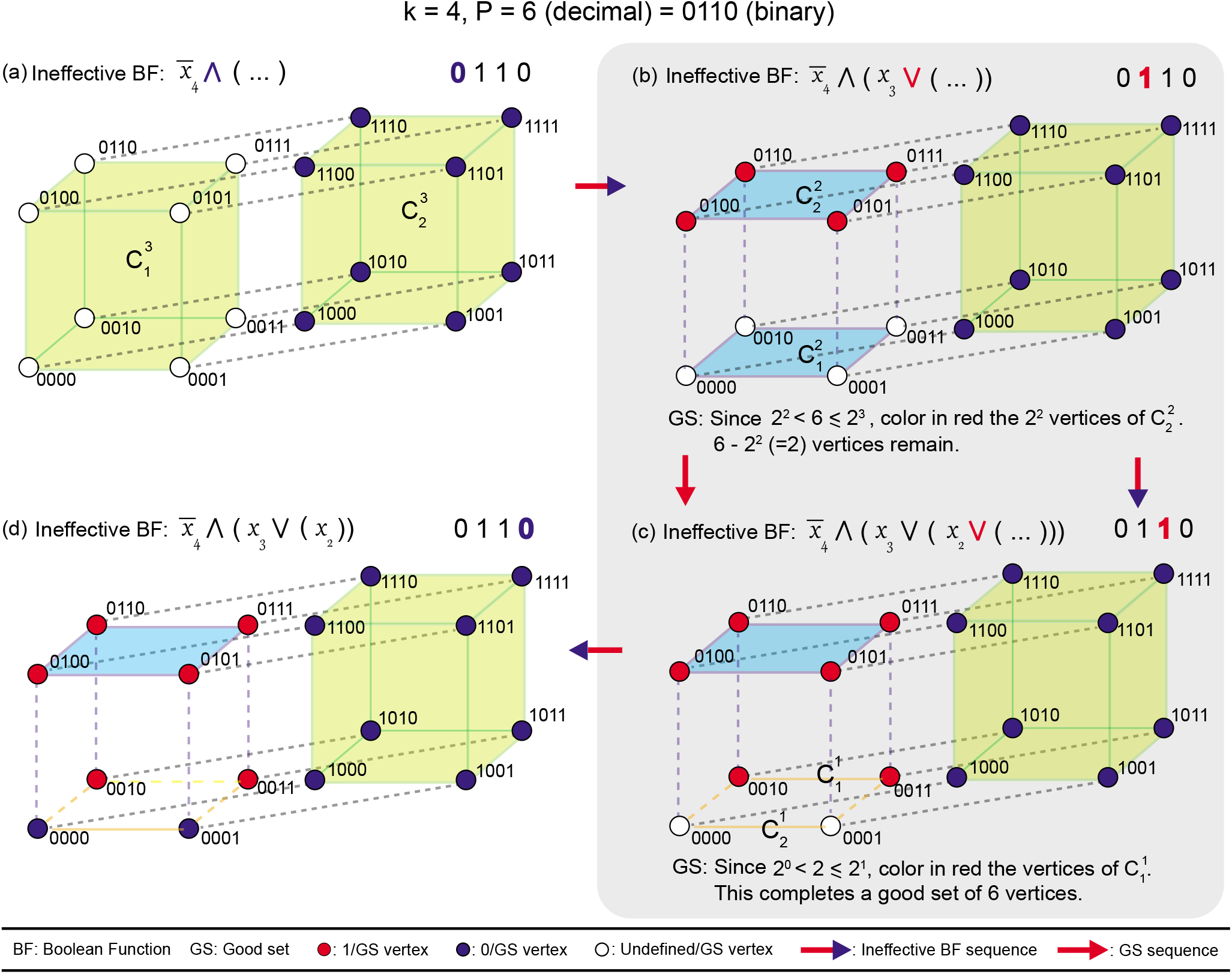
‘Good set’ (GS) for *P* vertices where *P* is even on a *k*-dimensional hypercube is equivalent to an ineffective Boolean Function (BF) in that *k*[*P*] set. In parts (b) and (c) shaded in grey, we show the recursive construction of a GS for *P* = 6 vertices in a 4-dimensional hypercube by coloring it’s vertices red, and in parts (a), (b), (c) and (d), we show the equivalence of that GS with 6 vertices to an ineffective BF with bias 6. The vertices of the hypercube are labeled in the order *x*_4_, *x*_3_, *x*_2_, *x*_1_ wherein *x_i_* is 0 or 1. Here, 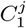 and 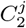 denote the two vertex disjoint *j*-dimensional hypercubes of the (j + 1)-dimensional hypercube. The ‘active’ bit in each part (a), (b), (c) and (d) is the colored bit in the binary representation of 6 in that part. **(a)** The vertices with *x*_4_ = 1 are set to the output value 0. The active bit in this step is 0 and as a result the ∧ operator follows the literal 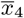. **(b)** Since *P* = 6 lies between 2^2^ and 2^3^, 2^2^ vertices of either 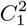 or 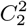 (here, 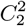) form part of the GS. This leaves 6 – 4 = 2 vertices to be colored to complete the GS. This choice of 4 vertices in 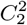 for the GS leads to setting the output value of vertices labeled *x*_4_ = 0 and *x*_3_ = 1 to 1. The active bit in this step is 1 and as a result the ∨ operator follows the literal *x*_3_. **(c)** Since the remaining 2 vertices lies between 2^1^ and 2^2^, 2^1^ vertices of either 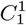 or 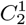 (here, 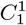) form part of the GS. This completes the GS of 6 vertices. This choice of 2 vertices in 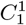 for the GS leads to setting the output value of vertices labeled *x*_4_ = 0, *x*_3_ = 0, and *x*_2_ = 1 to 1. The active bit in this step is 1 and as a result the ∨ operator follows the literal *x*_2_. **(d)** Since the two remaining undefined vertices take the same value 0, the variable *x*_1_ does not appear in the expression of the BF. Thus, the BF corresponding to GS with 6 vertices is ineffective.

**FIG. S7.**
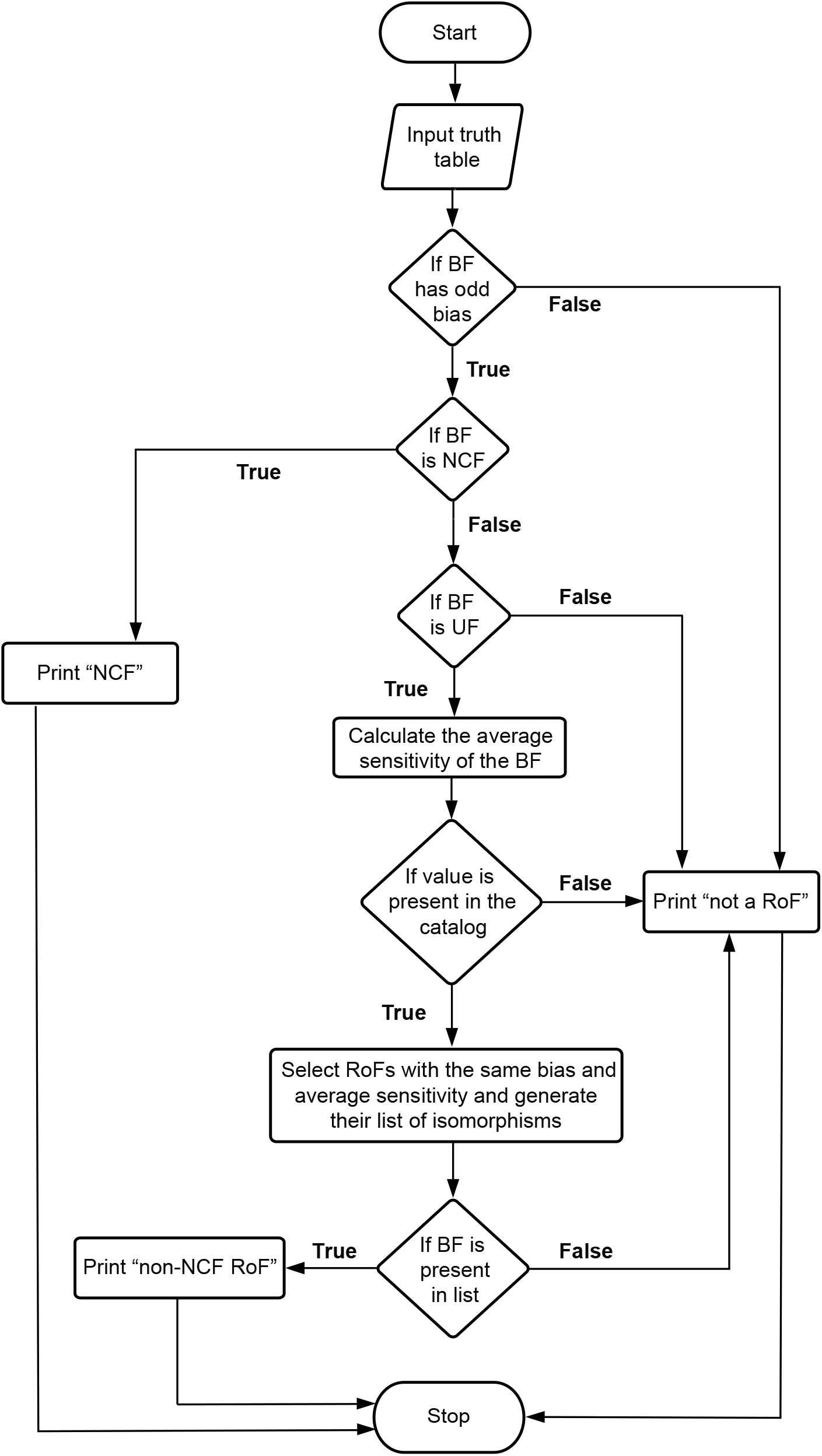
Flowchat describing our program to check whether a Boolean function (BF) with *k* ≤ 10 inputs entered by an user, is a Read-once function (RoF). The program can also distinguish between a Nested Canalyzing function (NCF) and a ‘non-NCF RoF’.

